# Modeling Sex Differences and Neurodegeneration in Repetitive Traumatic Brain Injury Using *Drosophila*

**DOI:** 10.64898/2026.02.27.708626

**Authors:** Nicole J. Katchur, Jessica Yeager, Lisa Schneper, Hacer Savas, Daniel A. Notterman

## Abstract

Traumatic brain injury (TBI) is a public health burden with short-term and long-term consequences for those who survive. Repetitive TBI (rTBI) is highly associated with neurodegenerative diseases such as chronic traumatic encephalopathy. Here, we present a force-adjustable, enhanced CO_2_-driven *Drosophila* model of rTBI to investigate locomotor, cognitive, proteomic, and neurodegenerative changes following rTBI. Injured males demonstrate dose-dependent locomotor deficits, while injured females display greater tolerance to rTBI. However, both sexes exhibit significant decision-making deficits. Proteomic analysis revealed changes in proteins linked with locomotory behavior, mitochondria, and energy production post-injury, correlating with observed behavioral phenotypes. Histological analysis revealed increased vacuolization area in rTBI flies. These results suggest that rTBI drives sex-specific behavioral deficits, proteomic disruption, and neurodegeneration. Overall, this enhanced *Drosophila* rTBI model provides a valuable tool for exploring sexually dimorphic outcomes and understanding the long-term pathophysiology of repetitive brain trauma in the context of neurodegenerative disease.

**Significance Statement:** Repetitive traumatic brain injury (rTBI) is linked to long-term cognitive decline and neurodegeneration, yet standardized, scalable models that capture sex-dependent outcomes are limited. We developed an automated, force-adjustable, closed-head *Drosophila* rTBI apparatus that delivers reproducible injuries without off-target damage, enabling precise control of injury severity across sexes. Using this platform, we show sexually dimorphic survival and locomotor vulnerability, persistent decision-making deficits, and progressive brain pathology marked by vacuolization and apoptosis. Proteomic profiling reveals time-dependent, sex-specific disruption of mitochondrial, oxidative stress, and locomotor-related protein networks, providing mechanistic entry points for discovering modifiers and therapies for repetitive brain trauma.

## Introduction

Traumatic brain injury (TBI) poses a significant public health risk, with long-term neurological and physical complications for those who survive. Recent evidence suggests a link between repetitive TBI (rTBI) and an increased risk of neurodegenerative diseases, including early-onset dementia and chronic traumatic encephalopathy (CTE), a progressive and often fatal condition associated with substantial exposure to repetitive head impacts and pathological deposition of aberrant tau (1, 2). TBI exposure lowers the age for cognitive decline onset, (3) and several studies suggest that biological sex is a significant risk factor for neurodegeneration (4–6). While rTBI is commonly associated with contact sports, it is also observed in other settings of head injuries (7) including military service (8), intimate partner violence (IPV), (9) and severe or uncontrolled seizures (10). Historically, TBI was thought to predominantly affect males (11), but over the last several decades, there has been a rise in TBI rates among females (12) due to increased military enrollment (13), participation in sports (14), IPV exposure (15), and greater awareness. The rise in TBI rates among females, the biological sex bias in research, and the variable short-term and long-term outcomes following injury underscore the need for a deeper understanding of how TBI outcomes are affected by sex differences. In recent decades, researchers have developed many animal models to evaluate TBI and its long-term consequences. Broadly, there are two main mechanisms of non-penetrating brain injury, inertial (acceleration/deceleration) and contact (16–18). To probe inertial mechanisms, Gennarelli and colleagues (1982) developed a diffuse axonal injury model in which the heads of nonhuman primates are accelerated in three directions without impact (19). To investigate contact TBI, other researchers have explored impact or contact models of injury, including a controlled cortical impact (CCI) device, wherein a pneumatic piston injures the exposed brain following a craniotomy (20–22).

Recent work has also included non-mammalian models of TBI in organisms such as nematodes and fruit flies. Using *Drosophila* to study single and repetitive TBI offers numerous potential benefits, including short longitudinal studies, straightforward genetic manipulation, and rapid throughput screening for potential therapies (23–26). In 2017, Sun and Chen developed a closed-head trauma model of CTE in *Drosophila* by using a ballistic impactor (27). However, that ballistic impactor presents several limitations, including calibration of the device to one sex and potential variability in the timing of the manual switch operation to shunt CO_2_ to the impactor (27).

Proteomic analysis is a powerful tool for understanding molecular processes through exploring complex networks, signaling pathways, co-expression, and post-translational modifications, allowing unbiased detection of sex-specific molecular signatures following injury. Researchers have used mass spectrometry, in animal models and humans, to elucidate potential biomarkers of TBI (28–33). Though one of these studies evaluated time-dependent molecular profiles of the brain after repetitive mild brain injuries in mice and in transgenic tauopathic models (expressing *Presenilin 1* mutations or human tau), the study did not include females (33). To our knowledge, no study has explored the proteomic changes associated with rTBI in both males and females. We develop and use an enhanced pneumatic ballistic impactor model of rTBI to investigate behavioral, proteomic, and neurodegenerative changes associated with repeated TBIs in both sexes of *Drosophila melanogaster*. We explore how repetitive brain injuries impact protein expression profiles in male and female flies, with a focus on identifying sexual dimorphism in response to injury and its correlation with behavioral outcomes.

## Results

### Injury variability and calibration challenges in a manual pneumatic rTBI model

To investigate the effects of repetitive traumatic brain injury (rTBI) in *D. melanogaster*, we initially used an existing design for a ballistic impactor system. The pneumatic apparatus developed by Sun and Chen was originally designed for female flies, delivering five impacts over five days, with a 24-hour recovery interval between each injury using a manual FlyBuddy regulator toggle to control the CO_2_ flow to the impactor (27). We define an impact or injury as a single closed-head TBI.

Sun and Chen demonstrated that a single impact delivered at a CO_2_ flow rate of 5.0 L/min produced a survival index of 100% after 24 hours (27). Therefore, we initially selected a 5.0 L/min flow rate as the median value for injury (range: 3.0 L/min–7.0 L/min). Male flies were subjected to five impacts at each flow rate (once per day for five days), and survival was monitored until death of all flies. In contrast to previous findings (27), rTBI at 5.0 L/min revealed significant differences in survival between flies with rTBI and control flies (χ² = 13.21, df = 1, p = 0.0003). The rTBI group had a median survival time of 7 days, compared to 42 days for the control group. There was a distinct separation between the survival distributions of the rTBI and control groups, with a greater proportion of rTBI flies dying earlier in the observation period. rTBI flies at 5.0 L/min had a five-fold increased risk of mortality compared to controls (HR = 5.2) (**Figure S1A**). Only 36.8% of male flies with rTBIs survived 24 hours after the final impact at a flow rate of 5.0 L/min (**Figure S1B**), indicating that 5.0 L/min leads to a rapid and severe decline in survival, limiting its use for longitudinal studies.

We then tested CO_2_ flow rates (3.0 L/min-7.0 L/min) to assess injury intensity effects on lifespan. Male flies exposed to 3.0 L/min showed no survival reduction (**Figure S1C**), with 80% surviving 24 hours post-final impact (**Figure S1D**). In contrast, male flies exposed to 5.0 L/min and 7.0 L/min had significantly decreased survival probability (χ² = 40.19, df = 3, p < 0.0001) (**Figure S1C**). Mortality risk was 3.6-fold higher at 5.0 L/min (HR = 3.63) and ten-fold higher at 7.0 L/min (HR = 10.1) compared to control flies. Furthermore, the survival index for flies exposed to rTBIs at 7.0 L/min was markedly reduced to 11.5% (**Figure S1D**). These results indicate that CO_2_ flow rates exceeding 5.0 L/min significantly reduce survival probability and the overall cohort size, thereby posing challenges to studying the longitudinal effects of rTBI.

While flipping the manual switch on and off, we observed a burst of CO_2_ flow in the regulator line that often exceeded the expected flow rate, suggesting that the regulator did not allow enough CO_2_ leak to stabilize the pressure in the CO_2_ line, resulting in inconsistent force delivered by the impactor. We also noticed that the time of positive pressure varied with the experimental trial and the operator. These observations suggested that the manual regulation of the CO_2_ flow introduced variability in the system, prompting us to equip the model with an automated, calibrated flow mechanism.

### Automated rTBI apparatus delivers reproducible injuries without off-target damage

We modified the Sun and Chen model by replacing the FlyBuddy regulator with a solenoid valve controlled by an Arduino Uno R3 for precise control of CO_2_ flow (**Figure S2A-B**). When initiated, the valve opens for 100 milliseconds (ms), directing a CO_2_ burst to accelerate the impactor to strike the *Drosophila* head. The valve closes after 100 ms, allowing the impactor to return to its starting position. An inline manometer allows precise titration and monitoring of the line’s CO_2_ pressure (pounds per square inch; psi) to promote reproducible brain injury.

Additionally, the valve opening time can be calibrated to the millisecond to ensure precision with each strike. To account for head size differences between Oregon-R wild-type *Drosophila* males and females, we modified 200 µL pipette tips, creating an 0.8-mm opening for males, and further enlarged for female flies without altering tip length. The kinetic energy of the ballistic impactor was calculated using the formula *KE = ½ * m * v^2^*, wherein *m r*epresents the mass of the impactor and *v* represents its velocity. The mass of the impactor was measured as 0.04 grams and its velocity (at 1 psi) was determined to be 0.96 m/s. Therefore, the kinetic energy of the impactor was 1.84 x 10^-5^ Joules when the pressure was set to 1 psi (**Figure S2C**). The calculated kinetic energy can inform the design and optimization of the ballistic impactor apparatus for various applications. Visual inspection of the *Drosophila* head following injury revealed no evidence of external damage to the cuticle, eyes, or proboscis (**Figure S2D**). This observation was consistent across multiple studies using the ballistic impactor device. Five rTBIs did not alter gut permeability (**Figure S2D**), suggesting no internal damage, as expected.

### Dose-dependent lifespan effects of CO₂ pressure levels in an automated rTBI apparatus

We assessed the reproducibility of the modified apparatus by observing the survival of female and male flies undergoing one injury daily for five days at varying CO_2_ pressures (1, 5, and 10 psi). Survival probability differed significantly between rTBI flies and control flies when the pressure exceeded 1 psi (males: χ² = 25.42, df = 1, p < 0.0001; females: χ² = 5.51, df = 1, p = 0.0189) (**Figure 1**). Male flies exposed to five TBIs at 5 psi or 10 psi demonstrated significantly reduced survival probability, with no difference at 1 psi (**Figure 1A**). In male flies, exposure to five injuries at 5 psi resulted in a 2.5-fold increased mortality risk (HR = 2.5); exposure to five impacts at 10 psi resulted in a further increased mortality risk (HR = 3.1) when compared to controls.

**Figure 1.**
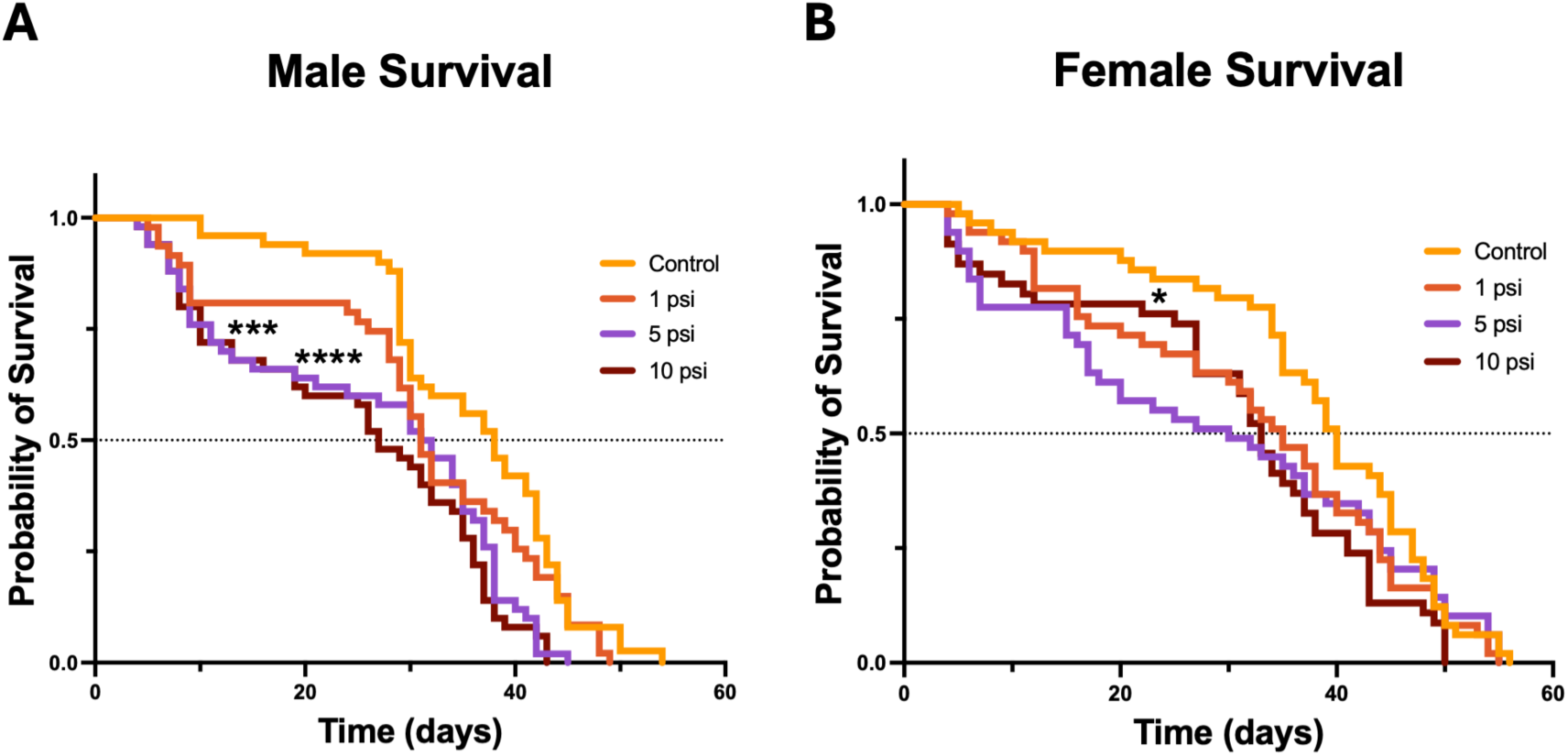
Effects of repetitive traumatic brain injuries on survival in male and female flies. Survival probability is shown for wild-type (*Oregon-R*) males (**A**) and females (**B**) subjected to five repetitive impacts at 1, 5, or 10 psi. No significant differences were observed at 1 psi in either sex. However, males exhibited significant reduction in survival probability at 5 and 10 psi, while females showed decreased survival probability only at 10 psi. Each group included 46–50 flies. Statistical significance: \**p* < 0.05, \*\**p* < 0.01, *\*\*\*\*p* < 0.0001; Log-rank trend test with Holm-Šídák’s multiple comparisons correction.

These results indicate a dose-dependent relationship between CO_2_ pressure (and strike energy) and mortality risk, with higher pressures leading to increased mortality risk. In female flies, five impacts at 10 psi increased mortality risk (HR = 1.8) when compared to controls (**Figure 1B**), while five impacts at 5 psi did not significantly decrease survival probability, suggesting males and females may differ in force tolerance following TBI. These results indicate that higher CO₂ pressures (and strike energy) correlate with greater injury severity and reduced survival. Survival probability in male and female controls did not significantly differ (**Figure S3A**), suggesting no sex-specific difference in overall lifespan in the absence of injury. Additionally, the difference between male and female survival probability at 1 psi was not statistically significant (**Figure S3B**), indicating that there is not a sex-specific difference in survival in response to repetitive injuries at 1 psi. These findings support the effectiveness of the modified apparatus in delivering reproducible and graded injuries to *Drosophila*, highlighting its potential as a model to study the acute and long-term consequences of rTBI, including sexual dimorphism.

### rTBI induces locomotor deficits in an automated rTBI apparatus

TBI may cause changes in cognition and physical ability, (34, 35) with gait and locomotor assays commonly utilized to evaluate TBI outcomes (36, 37). Male and female humans differ in symptom severity and long-term outcomes following rTBI (38, 39). Therefore, we evaluated locomotor performance in male and female flies one day post-injury (DPI). Male flies exposed to five injuries at 1 psi walked significantly shorter cumulative distances (2,173 mm ± 576.1) than age-matched controls (2,839 mm ± 665.5) (**Figure 2A-B**; U = 693, p < 0.0001), suggesting rTBIs produce significant walking deficits at one DPI in males. Female flies exposed to five injuries at 1 psi also walked significantly shorter cumulative distances (1,516 mm ± 385.7) than age-matched controls (2,037 mm ± 293.9) (**Figure 2C-D**; t(133) = 8.638, p < 0.0001). Two-way ANOVA revealed no significant sex-by-injury group interaction. These results indicate that the apparatus induces short-term locomotor deficits at 1 psi without evidence of sex-specific differences and are consistent with the survival findings at this setting.

**Figure 2.**
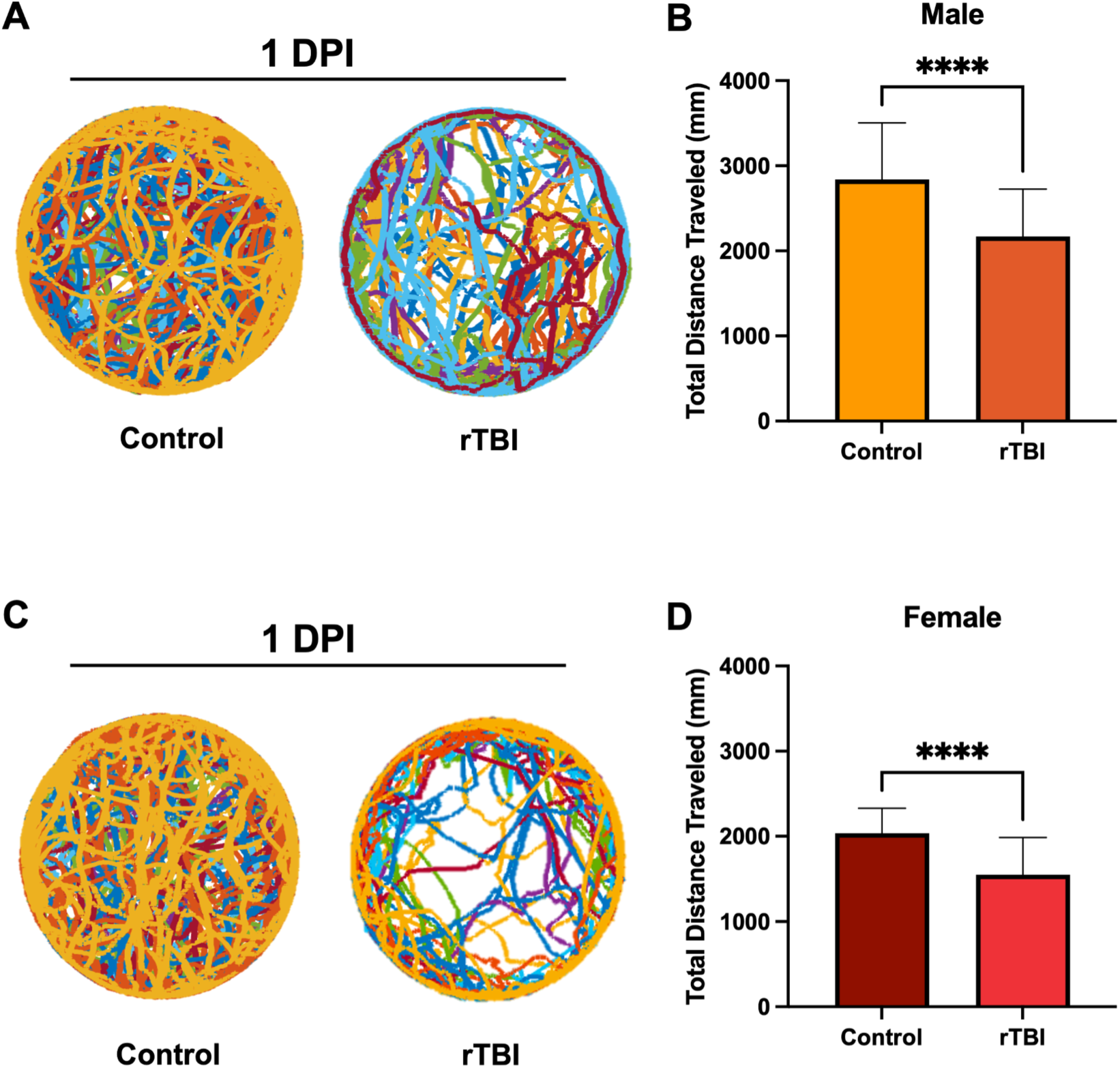
**Repetitive TBIs impair locomotor activity in *Drosophila.*** Representative 10-minute walking trajectories of wild-type (*Oregon-R*) males (A) and females (C) show reduced locomotor activity one day after repetitive TBIs (rTBIs; 1 psi). Cumulative walking distance was significantly decreased in males (B) and females (D) post-injury. Data are presented as mean ± SD. \**p* < 0.05, \*\*\**p* < 0.001, *\*\*\*\*p* < 0.0001; two-tailed Mann–Whitney U test. *n* = 20 flies per group. Two-way ANOVA revealed no significant interaction between sex and experimental groups, suggesting no sex difference in walking behavior following rTBI.

### Female flies exhibit greater tolerance to rTBI across injury thresholds

To explore long-term neurodegenerative sequelae following repeated head injury, we evaluated injury thresholds that produce behavioral changes while minimizing post-injury mortality. Locomotor deficits from TBI include slower walking speeds, abnormal gait, and increased support times(40, 41). Wild-type *D. melanogaster* flies were subjected to 1-5 head impacts with a 24-hour rest period between impacts, and locomotor behavior was assessed seven days after the final impact using the IowaFLITracker(42). In males, cumulative walking distance exhibited a non-linear relationship with injury number: males walked farther than controls after one TBI (3,389 mm ± 502.5, p < 0.0001 vs. control) and two TBIs (2,690 mm ± 382.9, p = 0.0299), but walked shorter distances after three TBIs (2,202 mm ± 983.9, p = 0.0009), four TBIs (2,488 mm ± 228.3, p < 0.0001), and five TBIs (2,470 mm ± 426.20, p = 0.0012) relative to controls (**Figure 3A**). In contrast to males, females showed a significant reduction in distance only after five TBIs (2,289 mm ± 304.6, p = 0.0192) (**Figure 3D**). Together, these findings suggest males are more sensitive to repeated head injuries with respect to locomotion, while females show greater resilience at lower injury numbers, with a deficit emerging only at the highest injury threshold tested.

**Figure 3.**
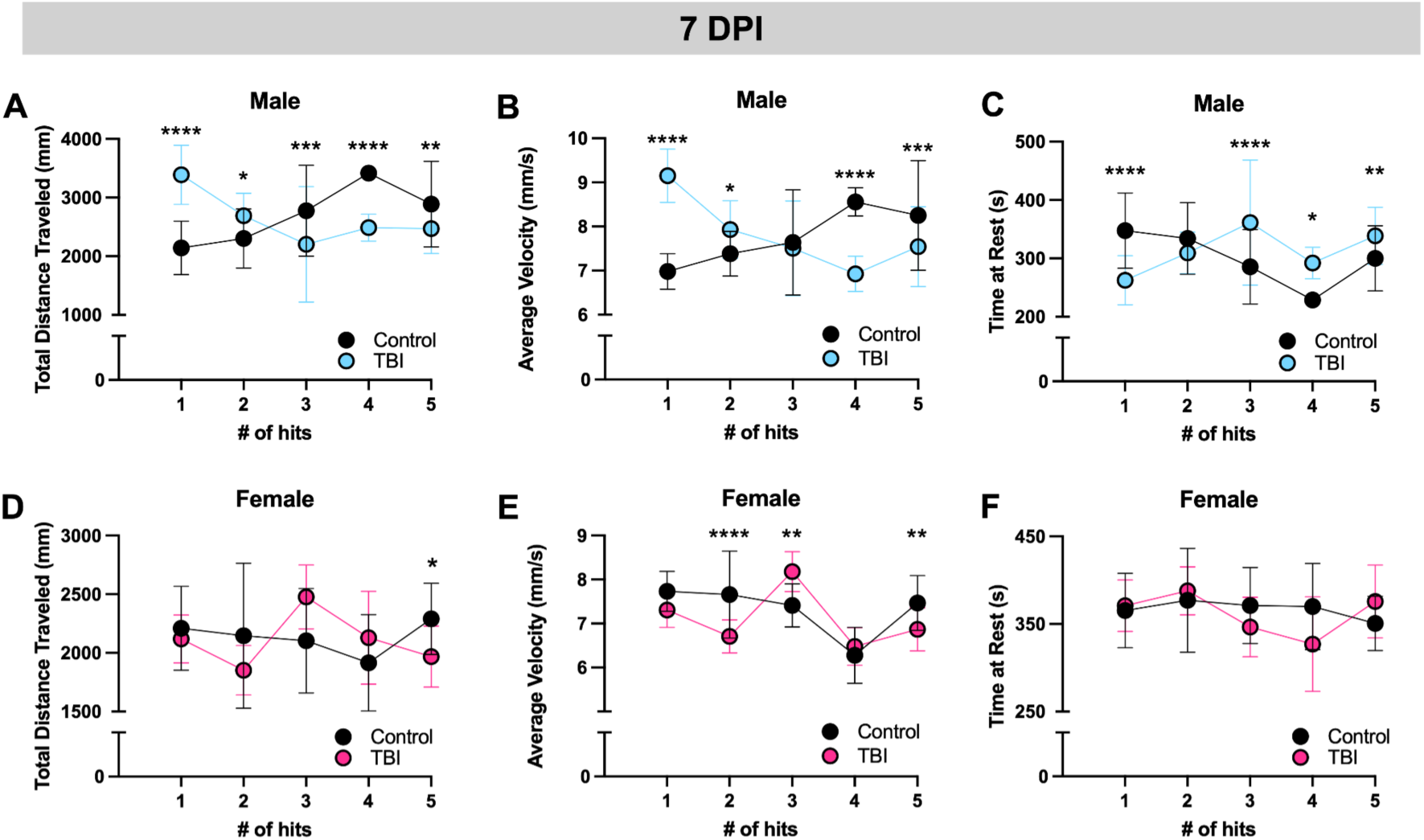
Repetitive TBIs induce persistent locomotor deficits in *Drosophila*. Flies were subjected to 1–5 TBIs at 24-hour intervals, and locomotor behavior was assessed seven days after the final injury. Total distance traveled (A, D), walking velocity (B, E), and rest periods (C, F) were quantified for males and females. Data are shown as mean ± SD. \**p* < 0.05, \*\**p* < 0.01, \*\*\**p* < 0.001, *\*\*\*\*p* < 0.0001; two-way ANOVA with Tukey’s multiple comparisons test. *n* = 11–40 flies per group.

To further characterize these impairments, we assessed the average walking speed of individual flies in five cohorts (1-5 injuries) and their respective age-matched control groups. Male flies displayed increased walking speed after one (9.15 mm/s ± 0.06, p < 0.0001) and two TBIs (7.93 mm/s ± 0.66, p = 0.0441) (**Figure 3B**), which aligned with the observed increase in total distance walked in these two cohorts. When male flies were exposed to four TBIs (6.9 mm/s ± 0.40, p <0.0001) and five TBIs (7.65 mm/s ± 0.58, p = 0.0002), the average velocity decreased when compared to age-matched controls, consistent with the reduction in distance traveled (**Figure 3B**). There was no change at three TBIs in males (**Figure 3B**). Analysis of female flies exposed to two (6.71 mm/s ± 0.38, p < 0.0001) and five TBIs (6.87 mm/s ± 0.49, p = 0.0018) revealed that injured females walked more slowly, while an increase in average velocity was observed in females exposed to three TBIs (8.18 mm/s ± 0.46, p = 0.0067) (**Figure 3E**).

Rest time (defined as movement below 3 mm/s) was also analyzed. Male flies rested significantly less after one TBI (262.6 s ± 42, p < 0.0001) but spent more time at rest following three (361.5 s ± 63.8, p < 0.0001), four (292.1 s ± 27.0, p = 0.0161), and five TBIs (338.5 s ± 49.0, p = 0.0057) (**Figure 3C**). These findings align with the observed increase with one TBI and subsequent reductions in walking distance and velocity. In contrast, female flies did not exhibit changes in their patterns of rest across all injury conditions (**Figure 3F**), suggesting that TBI primarily affects cumulative distance and average velocity in females.

Together, these results reveal injury-number-dependent locomotor deficits following rTBI, with males exhibiting greater sensitivity to repeated impacts as compared to female flies. Both sexes demonstrated significant impairments after five TBIs, establishing this injury threshold as a potential model for investigating long-term neurodegenerative outcomes associated with repeated head trauma.

### Defects in locomotion and cognition persist after rTBI for at least one week after injury

To better define the long-term effects of rTBI on locomotor behavior, we exposed male and female *Drosophila* to five head impacts and assessed walking behavior using IowaFLITracker software at multiple time points post-injury. One day after injury, males walked 2,173 mm ± 576.1 compared to the control distance of 2,839 mm ± 665.5 (p < 0.0001) and females walked 1,516 mm ± 385.7 compared to control distance of 2,037 mm ± 293.9 (p < 0.0001). This deficit persisted at seven DPI in both sexes, with injured males and injured females continuing to walk shorter distances than controls (males: 2,470 mm ± 426.2, p = 0.0156; females: 1,968 mm ± 260.7, p < 0.0001) (**Figure 4A,C**). However, females still exhibited walking deficits at 14 DPI (1,719 mm ± 478, p = 0.023) when compared to controls, but males did not. By day 21, injured males and female flies recovered. These findings suggest that females take longer to recover after repetitive head trauma than males, indicating a sex-specific difference in recovery following rTBI. At 14 days post-injury, after normalizing each fly’s locomotor distance to the sex-matched control mean at the same time point, males walked greater distances than females (male: 110.9% of control; female: 85.3% of control; mean difference = 25.6%, Welch’s t(62.4) = 3.91, p = 0.0002; Holm-Bonferroni adjusted p = 0.0009; Hedges’ g = 0.91 (large)). No sex differences were detected at 1, 7, or 21 days post-injury after the same normalization.

**Figure 4.**
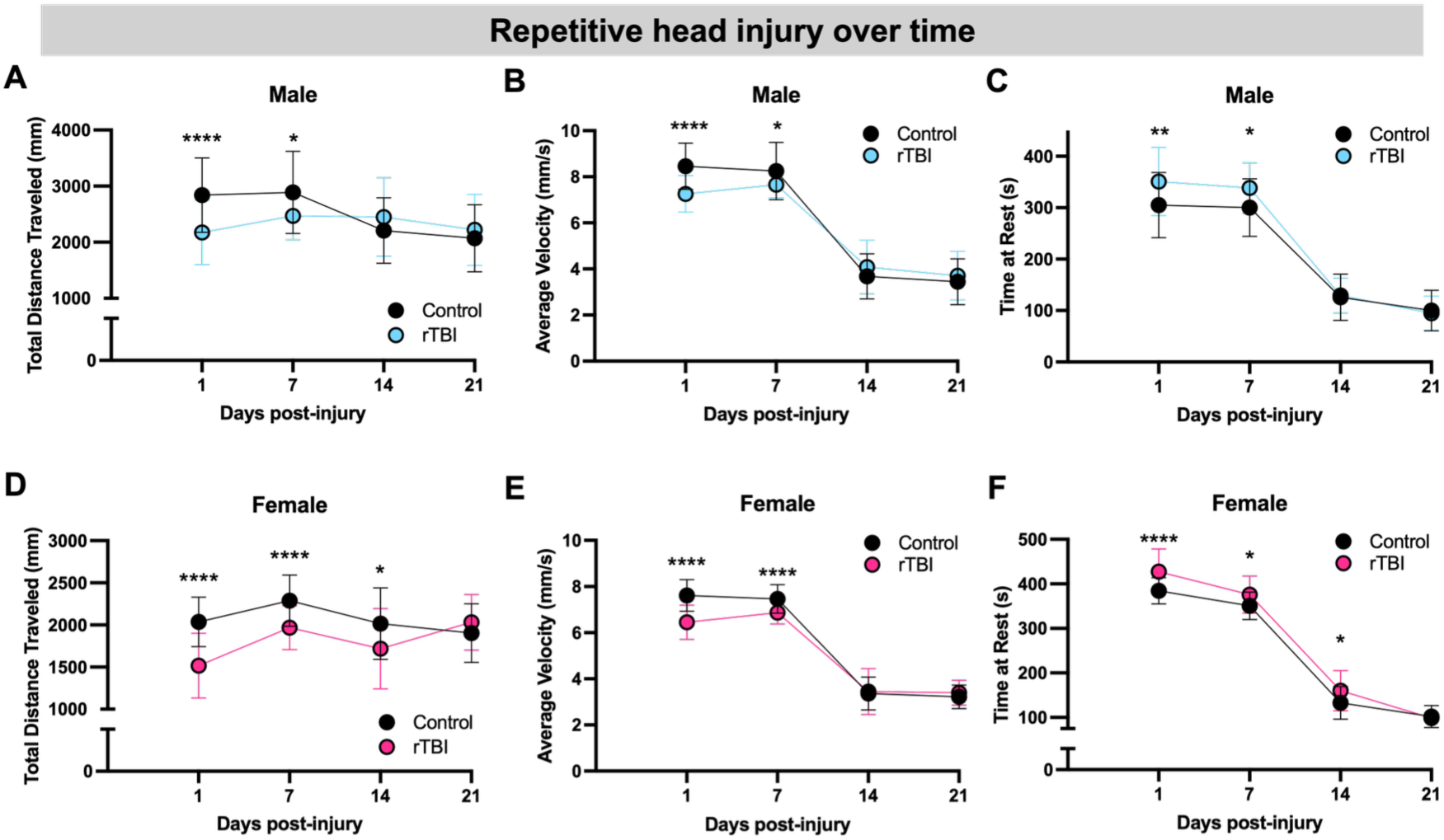
Longitudinal analysis of locomotor performance following rTBI. Cumulative distance walked over 10 minutes (A, D), mean velocity (B, E), and total rest time (C, F) were measured in males and females at 1, 7, 14, and 21 days post-injury. Injured flies displayed sustained locomotor deficits relative to age-matched controls. Data are presented as mean ± SD. \**p* < 0.05, \*\**p* < 0.01, *\*\*\*\*p* < 0.0001; two-way ANOVA with Tukey’s multiple comparisons test. *n* = 30–75 flies per group.

Across all time points, a factorial model including Sex, Injury, and Time revealed a significant three-way interaction (Sex × Injury × Time: *F*= 4.1, *p* = 0.0067), indicating that the effect of injury varied across time differently for males and females. Post hoc, multiplicity-adjusted contrasts localized this effect to day 14, when males and females diverged in their response to injury. Interestingly, control male and female flies demonstrated a significant difference in walking behavior, with control males exhibiting higher levels of locomotor activity compared to control females at each time-point (p < 0.0001). Control males may have greater cumulative walking distances because they engage in courtship behaviors such as chasing (43).

Older flies walk more slowly (44), and individuals with TBIs experience gait abnormalities, including a slower walking speed when compared to individuals without TBI (40, 45). Therefore, we expected our control flies to decrease in average velocity as they age, but we expected injured flies to have greater deficits in velocity when compared to normally aged controls. As expected, control male and female flies walked more slowly as they aged. Injured males and females both walked significantly slower one (males: 7.25 mm/s ± 0.80, p < 0.0001; females: 6.46 mm/s ± 0.74, p < 0.0001) and seven DPI (males: 7.65 mm/s ± 0.58, p = 0.0412; females: 6.87 mm/s ± 0.50, p < 0.0001) (**Figure 4B,E**). Interestingly, these significant differences were resolved by 14 DPI.

At early time points (one and seven DPI), when injured flies exhibited reductions in both cumulative distance and velocity, we expected a concomitant increase in rest time. Indeed, injured males spent significantly more time at rest one (350.7 s ± 66.3, p = 0.0015) and seven days (338.5 s ± 49, p = 0.0104) post-injury when compared to age-matched controls (**Figure 4C**). Similarly, injured females spent significantly more time at rest one (427.3 s ±51.2, p < 0.0001), seven (375.7 s ± 41.6, p = 0.0179), and 14 DPI (160.3 s ± 45.3, p = 0.0213) when compared to controls (**Figure 4F**). At 14 DPI, females continued to show reduced distance despite resolution of velocity differences, suggesting that persistent increases in rest time could underlie the distance deficit. Although injured flies rested more than age-matched controls at early time points, rest time decreased with age across the assay window in both injured and control cohorts (**Figure 4C,F**). This pattern contrasts prior reports that aging flies become less active (44, 46, 47).

### Short-term cognitive deficits persist in both sexes following rTBI

Persistent cognitive deficits following TBI are more common in female humans, manifesting as impairments in attention, decision-making, problem-solving, and judgment (48–51). To evaluate decision-making deficits following rTBI, we utilized the value-based feeding decision (VBFD) paradigm (52). In wild-type *Drosophila*, VBFD remains intact with age until midlife (52). *Drosophila* were forced to choose between a nutritious sucrose solution and a non-nutritious arabinose solution (**Figure 5A**). Healthy flies preferentially consume sucrose.

**Figure 5.**
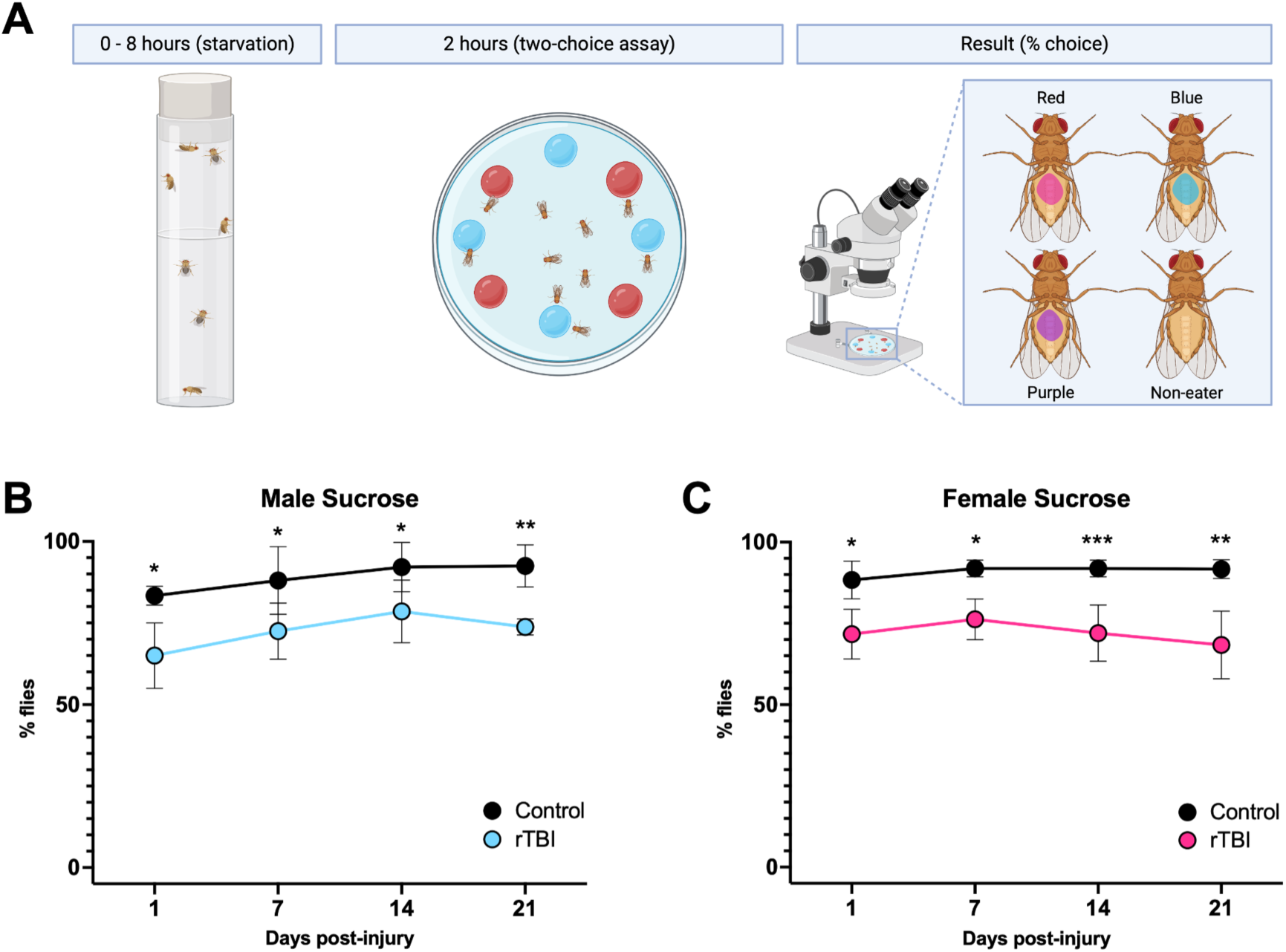
Repetitive TBIs impair cognitive function in *Drosophila*. (A) Schematic of the feeding preference assay where flies choose between sucrose (nutritious) and arabinose (non-nutritious) solutions. Feeding preference was quantified in males (B) and females (C), revealing short- and long-term impairments post-rTBI. Data are shown as mean ± SD. \**p* < 0.05, \*\**p* < 0.01, \*\*\**p* < 0.001; two-way ANOVA with Tukey’s multiple comparisons test. *n* = 10 flies per group; 3–7 groups per time point.

Percentages were calculated based on the number of flies that chose nutritious sugar over the total number of flies tested, including non-choosers. Male and female control flies exhibited a strong preference for sucrose regardless of age. However, injured males and females displayed significant deficits at all time points: 1 DPI (males: 65% ± 10, p = 0.0170; females: 71.7% ± 7.6, p = 0.0351), 7 DPI (males: 72.5% ± 8.7, p = 0.0133; females: 76.3% ± 6.3, p = 0.0123), 14 DPI (males: 78.5% ± 9.6, p = 0.0128; females: 72.0% ± 8.7, p = 0.0001), and 21 DPI (males: 73.8% ± 2.5, p = 0.0035; females: 68.3% ± 10.4, p = 0.0015) (**Figure 5B, C**). Significant differences between arabinose and sucrose consumption were observed in females at all time points (**Figure S4B**), but not at one and 14 DPI in males (**Figure S4A**), possibly due to the proportion of flies that did not choose a solution (**Figure S4C-D**). These findings suggest that rTBI impairs decision-making in *Drosophila*, but deficits are not observed in both sexes.

### rTBI induces significant neurodegeneration 14 days post-injury

Neurodegeneration, characterized by abnormally enlarged cell bodies, apoptosis, and vacuolization, is a hallmark of tauopathies, including CTE (53). To study rTBI-induced neurodegeneration, we analyzed male fly brains 14 DPI. Flies with rTBI had significantly greater vacuolization (0.36% of total brain section area per fly) than controls (0.14%; t(4) = 4.033, p = 0.0157) (**Figure 6A**). Enlarged cells on the brain periphery suggested cellular pathology consistent with apoptosis (**Figure 6C**), confirmed by TUNEL staining, which revealed apoptotic cells along the eye and optic medulla border (**Figure 6D**).

**Figure 6.**
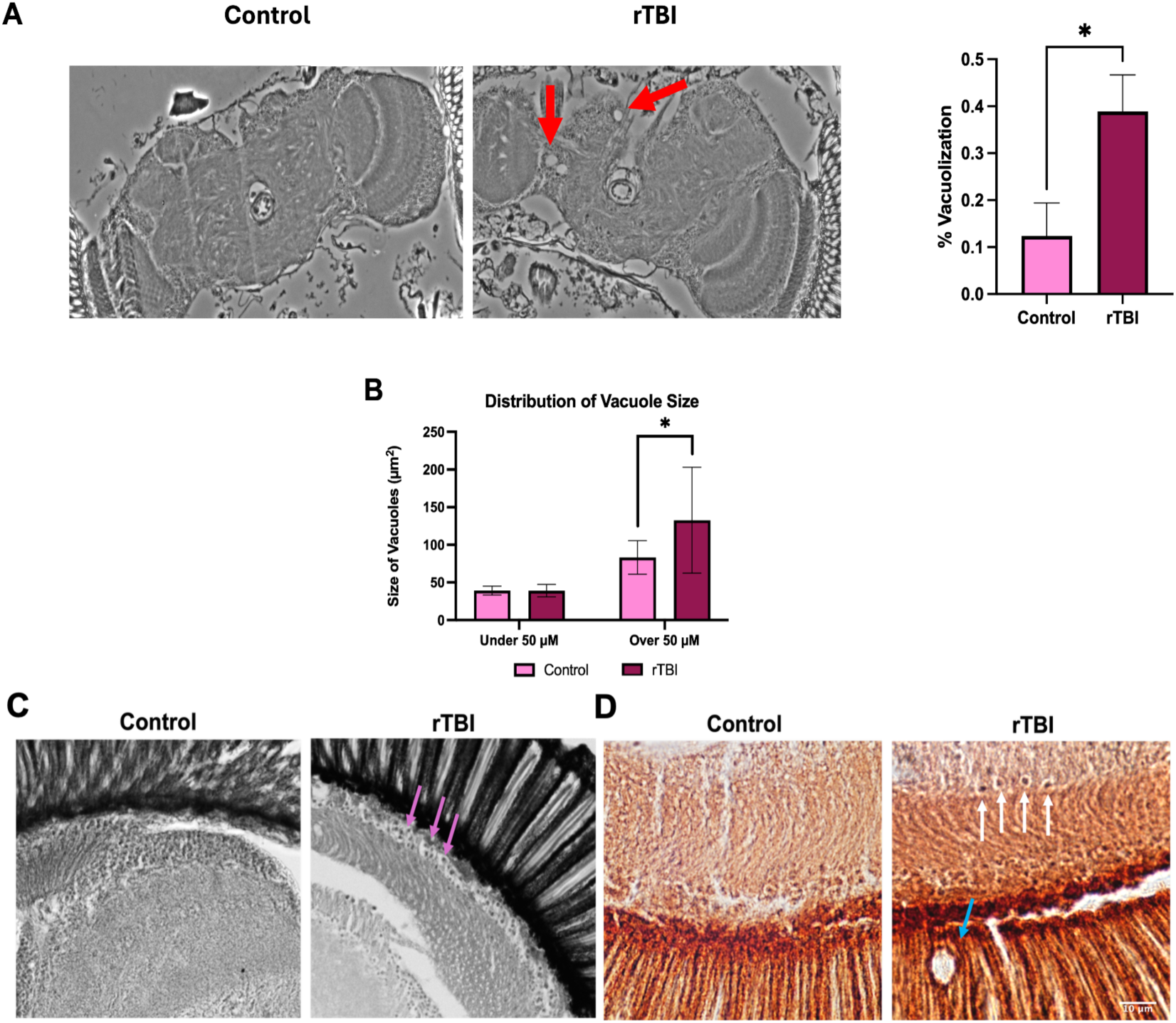
rTBI induces neurodegeneration and apoptosis in the *Drosophila* brain. (A) Representative images showing vacuolization (>30 µm², red arrows) in the optic medulla of injured wild-type flies. Quantification reveals increased brain tissue damage after rTBI. (B) Distribution of vacuole sizes: small (<50 µm²) and large (>50 µm²). (C) H&E-stained brain sections reveal enlarged cells (magenta arrows) indicative of apoptosis 14 days post-injury. (D) TUNEL staining highlights apoptotic cells (white arrows) near the medulla and eye 1 day post-injury; vacuolization is also observed (blue arrow). Data are shown as mean ± SD. \**p* < 0.05; unpaired two-tailed t-test. *n* = 3 sections per brain, 3 brains per group.

Vacuoles were categorized into small (<50 µm²) and large (>50 µm²) to differentiate age-related degeneration from injury-induced changes (54). Control and rTBI groups had similar numbers of small vacuoles and the average area of vacuolization of the small vacuoles was not statistically significant, indicating that these smaller vacuoles may be a result of normal aging (28 days post-eclosion) (**Figure 6B**). However, the rTBI group exhibited an increased number of large vacuoles compared to controls and the mean area of vacuolization was significantly larger in rTBI flies than controls (rTBI: 132.7 µm^2^ ± 70.2, control: 83.28 µm^2^ ± 22.3, p = 0.0385) (**Figure 6B**), suggesting that rTBIs induce the formation of larger vacuoles and contribute to an overall increase in total vacuolization following injury. We next assessed the regional distribution of vacuoles across brain subregions (see Supplemental Material).

### rTBI induces significant vacuolization in dTauKO flies

While tau has been linked to neurodegeneration and neurotoxicity, there is only a rudimentary understanding of the upstream biochemical mediators of tau in the context of rTBI and CTE. Several studies have expressed wild-type and mutant human tau (hTau) proteins in *D. melanogaster* to model Alzheimer’s disease (AD). However, hTau transgene expression in *Drosophila* is not an ideal functional model, in part because of poor binding to *Drosophila* microtubules. This, as well as differences in phosphorylation sites and uncertainty about whether hTau expression recapitulates endogenous neurofibrillary tangles (NFT) formation in flies, limits the applicability of this model. For example, *Drosophila* tau (dTau) contains five putative microtubule-binding repeats and lacks the N-terminal repeats that exist in human tau, despite sharing 66% homology with hTau protein. At least six CTE-associated phosphorylation sites are observed in human tau, and four of those, Thr^231,^ Ser^202^, Thr^205^, and Ser^199^, are conserved in dTau, as Thr^151^, Ser^106^, Thr^123^, and Ser^103^, respectively, when aligned using UniProt (**Figure S7**). Therefore, pathologic changes associated with expression or overexpression of hTau may not reflect the actual contribution of dTau to neurodegeneration in fly-based models of CTE or related disorders. Accordingly, we sought to utilize a *Drosophila* tau (dTau) knockout (KO) model to investigate neurodegeneration in the presence and absence of dTau after rTBI. After receiving successive strikes over five days, there was a significant difference in percent vacuolization between dTauKO injury and dTauKO control groups (**Figure 7A**), suggesting that neurodegeneration after rTBI is not strictly dependent on dTau. While rTBI wild-type flies showed tau accumulation around vacuoles at 14 DPI and 28 DPI, dTauKO flies did not show this accumulation (**Figure 7B**).

**Figure 7.**
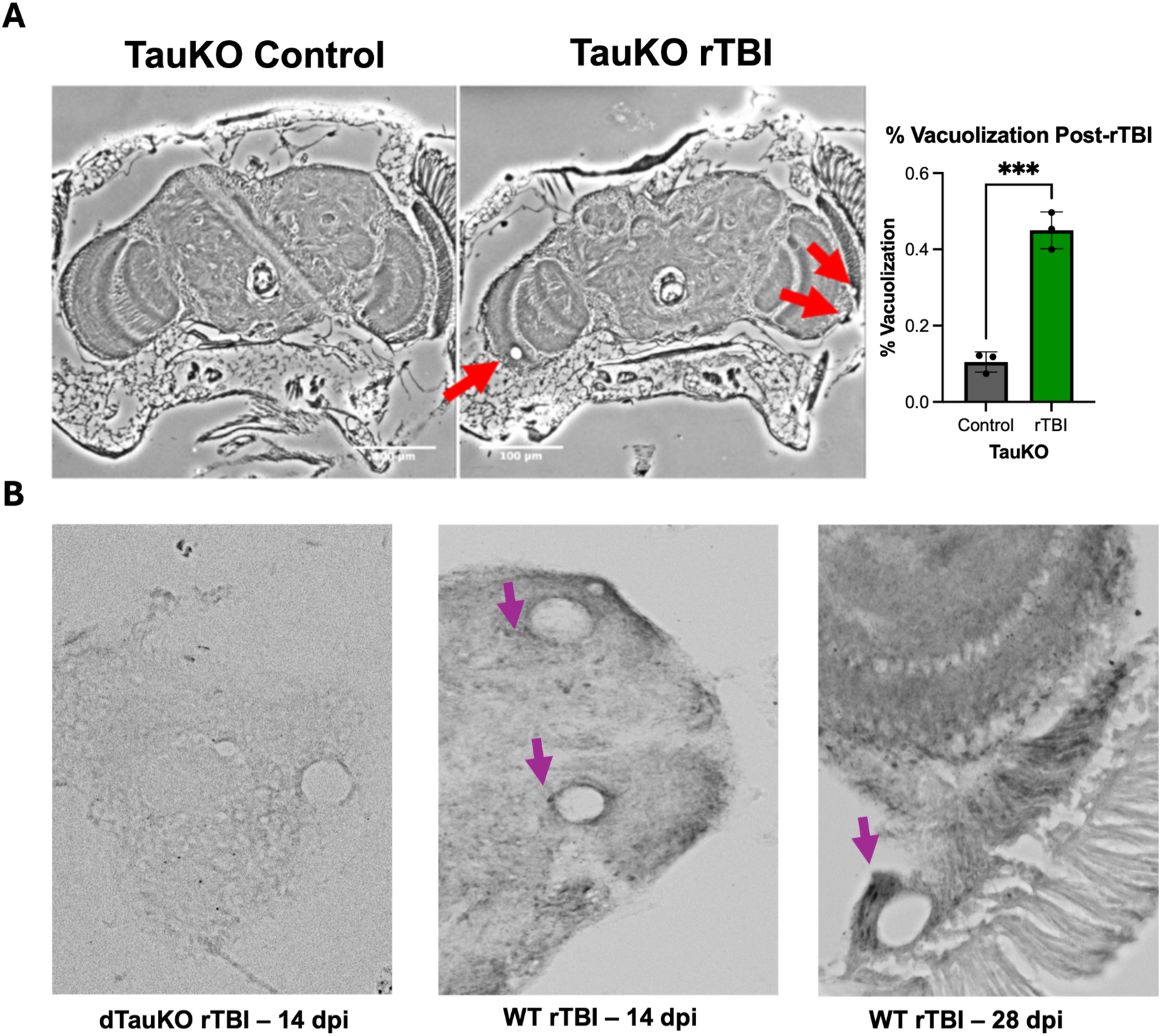
rTBI promotes vacuolization in dTauKO flies. (A) Representative images showing vacuolization (>30 µm², red arrows) in the optic medulla of dTauKO flies post-injury (left). Quantification indicates significantly increased vacuolization in injured dTauKO flies relative to controls (right). (B) Histological images show dTau accumulation (purple arrows) around vacuoles in wild-type flies but not in dTauKO flies. Data are presented as mean ± SEM. \*\*\**p* < 0.001; unpaired two-tailed t-test. *n* = 3 flies per group.

### Proteins linked to behavior, supramolecular fiber organization, and energy production altered by progressive injuries

Since both single and repetitive traumatic brain injuries have been associated with CTE-like phenotypes, the accumulation of kinetic energy, not the number of injuries, may account for the transition from TBI sequelae to neurodegenerative disease. To investigate the effects of accumulating injury load, we performed shotgun proteomics on male flies subjected to single or multiple head impacts. Proteomic analysis identified 743 unique proteins. Hierarchical clustering revealed an abrupt change in protein abundance pattern after three injuries (clusters 1-5), suggesting three impacts (*KE = 5.53 x 10^-5^ Joules)* may be a critical threshold for brain injury tolerance in this model (**Figure 8A)**. Gene ontology (GO) enrichment analysis revealed significant enrichment of terms related to homeostasis regulation, energy production, mitochondria, locomotor behavior, and supramolecular fiber organization (**Figure 8B**). Key proteins associated with locomotion (e.g., cabeza (caz), Akap200, Lam, and SERCA) and mitochondrial function were altered (**Figure 8B).** Many terms were linked to mitochondria and metabolism, suggesting the brain was either adapting to altered energy dynamics or became dysregulated in response to accumulating inflicted energy. These findings underscore the progressive impact of TBIs on brain protein networks.

**Figure 8.**
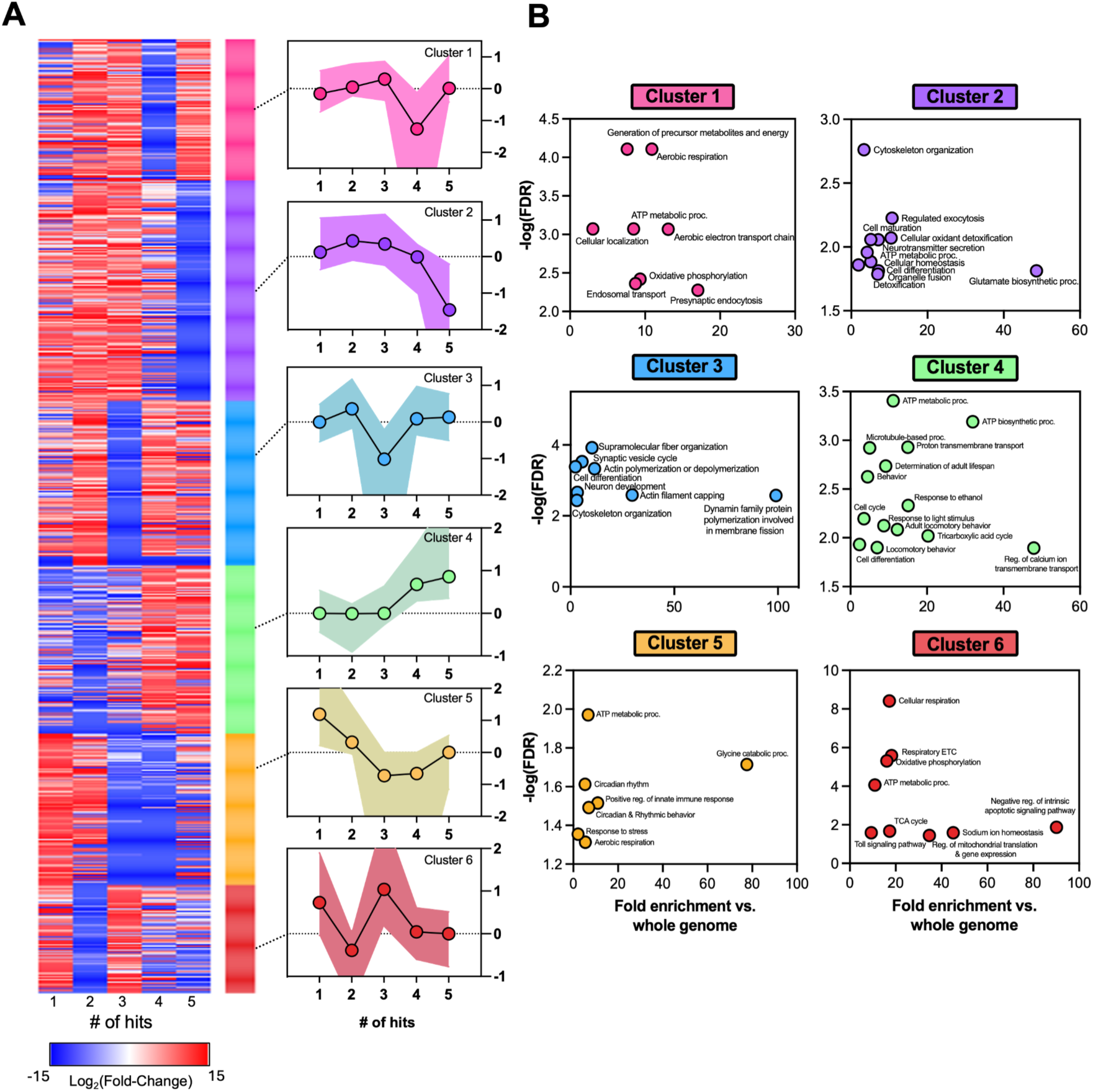
Proteomic profiling reveals injury-dependent molecular changes in male flies. (A) Mass spectrometry identified 743 proteins altered by 1–5 TBIs. Heatmap depicts fold changes in protein abundance across injury conditions. K-means clustering (k = 6; Morpheus) grouped proteins with similar expression patterns; median and interquartile ranges are shown. (B) Gene ontology (GO) analysis highlights enriched biological processes (ShinyGO 0.80; fold enrichment and FDR values reported; https://bioinformatics.sdstate.edu/go/).

### Longitudinal proteomic changes in male and female flies post-rTBI

To investigate sex-linked changes in protein abundance after rTBI across recovery, we exposed male and female flies to five head impacts and quantified proteomic changes at 1, 7, and 14 DPI. The dataset comprised 2,602 proteins mapped to the *Drosophila* proteome. Hierarchical *k*-means clustering of injury-associated log_2_ fold-changes identified six protein clusters (Clusters 1–6) with distinct sex- and time-dependent trajectories (**Figure 9A**).

**Figure 9.**
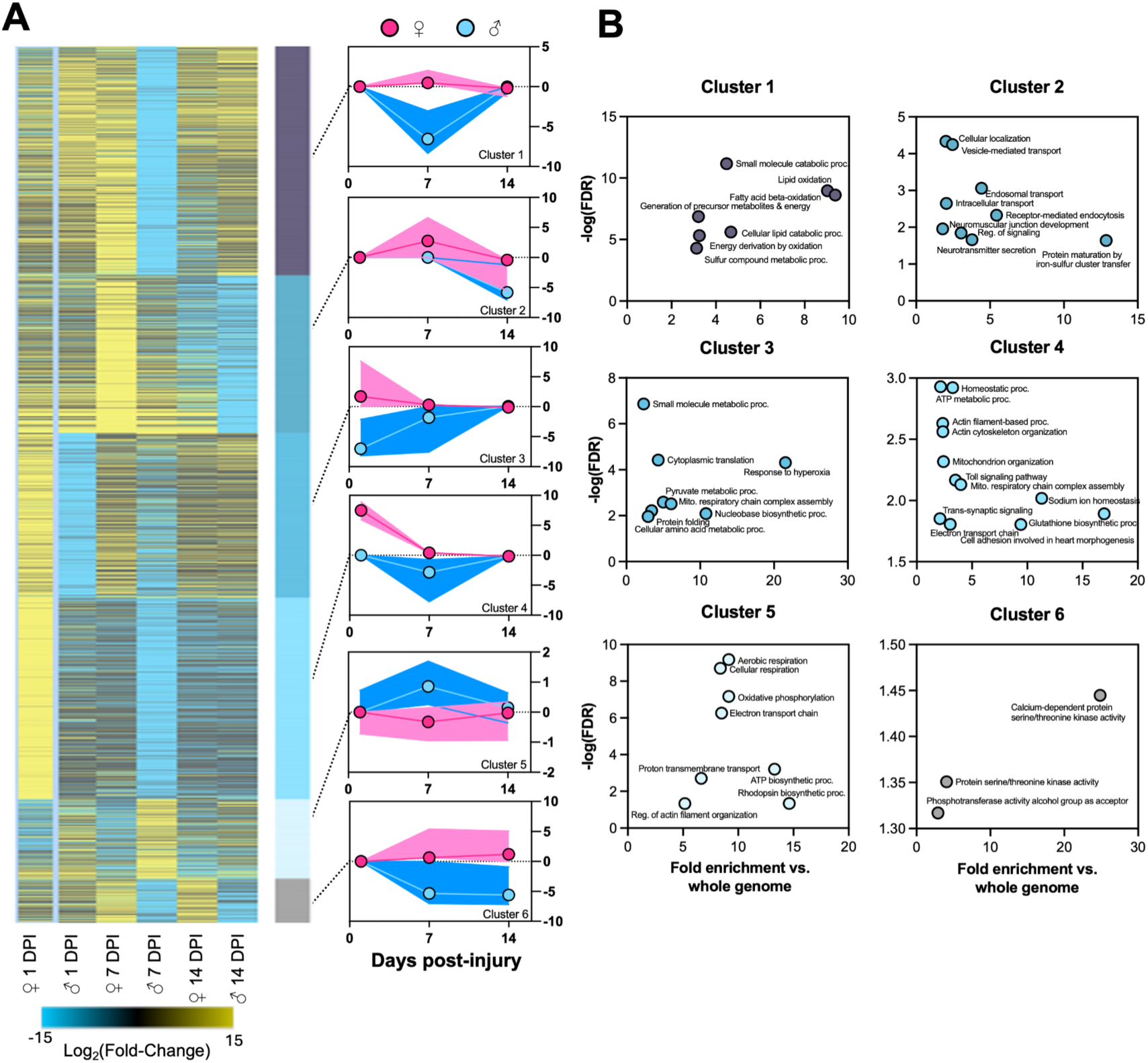
Longitudinal proteomic changes in male and female flies post-rTBI. (A) Mass spectrometry identified 2,602 proteins in injured flies at multiple time points. Heatmap shows relative abundance changes in males and females. K-means clustering (k = 6; Morpheus) identified co-regulated protein groups; median and interquartile ranges are plotted for each cluster. (B) GO analysis highlights injury-associated pathways (ShinyGO 0.80; fold enrichment and FDR values reported).

Several clusters showed the largest divergence at 7 DPI, particularly in males. Cluster 1, enriched for lipid oxidation and energy derivation pathways (fatty acid β-oxidation and generation of precursor metabolites), exhibited a pronounced decrease in males at 7 DPI with recovery toward baseline by 14 DPI, whereas females remained near baseline across time (**Figure 9A,B**). Cluster 4, enriched for homeostatic processes, mitochondrial organization/respiratory chain assembly, cytoskeletal organization, and Toll signaling, showed a male-biased reduction at 7 DPI with recovery by 14 DPI; females showed the largest deviation at 1 DPI (**Figure 9A,B**). In contrast, Cluster 5, enriched for aerobic respiration, oxidative phosphorylation, and electron transport chain terms, displayed higher abundance in males at 7 DPI with partial resolution by 14 DPI, while females showed a comparatively muted trajectory (**Figure 9A,B**). Together, these patterns indicate that mitochondrial and metabolic networks are dynamically remodeled after rTBI, with the most prominent shifts occurring at 7 DPI and exhibiting sex specific differences in magnitude and direction (**Figure 9**).

Other clusters exhibited sex divergence at specific time points rather than a shared peak at 7 DPI. Cluster 2, enriched for vesicle-mediated transport, endosomal trafficking, neuromuscular junction development, and neurotransmitter secretion, increased in females at 7 DPI and returned toward baseline by 14 DPI, whereas males showed a sharp decrease at 14 DPI (**Figure 9A,B**).

Cluster 3, enriched for cytoplasmic translation, small-molecule metabolism, mitochondrial respiratory chain complex assembly, and response to hyperoxia, showed opposing sex patterns at 1 DPI with convergence by 14 DPI (**Figure 9A,B**). Finally, Cluster 6, enriched for calcium-dependent serine/threonine kinase activity (including CaMKII), exhibited the most persistent sex divergence: females increased at 7 and 14 DPI, whereas males decreased at both time points (**Figure 9A,B**), consistent with sustained sex-specific perturbation of calcium signaling after injury.

To contextualize these cluster trends, we examined sex-by-time fold-changes for individual proteins after manually clustering the data (**Figure S6**). At 14 DPI, several annotated proteins displayed sex-biased regulation (e.g., caz increased in males while decreasing in females, whereas amon and sxc increased in females despite minimal or opposite changes in males; **Figure S6C**). Collectively, these results indicate that rTBI perturbs mitochondrial/metabolic, trafficking/synaptic, and calcium-dependent kinase pathways in a sex-and time-dependent manner, with a prominent transition at 7 DPI and persistent divergence in calcium/kinase-associated proteins through 14 DPI.

## Discussion

The present study links repetitive traumatic brain injury to longitudinal behavioral, proteomic, and histologic outcomes in *Drosophila*, with several measures exhibiting sex-dependent recovery dynamics. Given that TBI has both acute and chronic consequences often leading to lasting cognitive, neuroendocrine, and physical stigmata (34, 35), and is associated with long-term risk for neurodegenerative disease, including AD and CTE (55–57), model systems that permit systematic manipulation of injury parameters and long-term follow-up are essential for identifying mechanisms that may underlie divergent outcomes across individuals and sexes. A major contribution of this work is the development of a more reproducible head-injury apparatus suitable for both male and female flies. Prior *Drosophila* TBI models can be variable in intensity or lack head specificity (23, 24), and the Sun and Chen ballistic impactor improved this space by delivering head-specific impacts to non-immobilized flies(58), thereby approximating relevant concussion dynamics (59). However, manual operation, timing variability, and design for a single sex limit reproducibility and generalizability (58). By digitally monitoring pressure and automating control valve timing, our modified device enables controlled injury delivery in both males and females and supports longitudinal behavioral and molecular profiling.

Using this apparatus, we identified an injury protocol that models mild rTBI: five head impacts at 1 psi (*KE = 9.22 x 10^-5^ J*) produced robust short-term locomotor deficits in both sexes without altering survival, consistent with mild concussion-like phenotypes (60). Importantly, recovery trajectories diverged by sex. While both sexes were impaired early, males recovered locomotor distance by 14 DPI, whereas females exhibited a more prolonged deficit extending to 21 DPI. This aligns with clinical observations that females experience higher risk of poor outcomes months after mild TBI despite similar rates of return to daily activities (38). Women with mild TBI often experienced longer delays to CT scans and were more likely to be discharged home (61), while reporting more post-concussive symptoms than men after three months (38). As TBI incidence among females rises due to sports, military service, and IPV, and females remain underrepresented in many TBI studies, defining sex-specific recovery patterns in tractable models remains critical for reducing bias in mechanistic understanding and therapeutic development.

In the broader context, animal studies investigating rTBI vary widely in terms of time intervals between repeat head injuries and the number of head injuries administered (58, 62, 63), highlighting the lack of a standardized approach for inducing rTBIs. Variability may lead to differences in observed outcomes and make it challenging to compare between model systems. To address this directly, we systematically varied injury number while holding inter-injury interval constant (24 hours) and assessed behavior at 7 DPI. In males, locomotor output showed a non-linear relationship with injury number, with early increases in distance and speed at 1-2 injuries followed by impairment starting at three injuries (*KE* = 5.53 x 10^-5^ J), whereas females exhibited a significant distance reduction primarily at the highest injury threshold tested. Together, these findings support sex differences in injury tolerance or compensatory responses where females appear comparatively resilient to lower numbers of strikes but show persistent impairment once a threshold is crossed. The literature reports that children are often more vulnerable to the force dynamics of a TBI because of their relatively larger head-to-body ratio when compared to adults (64). Although female flies have larger heads than males, their head and body dimensions scale proportionally, and the apparatus was calibrated to accommodate sex-specific differences in head size. Accordingly, gross head size alone is unlikely to explain the observed sex-dependent injury tolerance. Instead, sex-dependent molecular responses likely contribute, consistent with the sexually dimorphic proteomic trajectories observed after injury.

A central implication of our proteomic analyses is that cumulative force, rather than injury number *per se*, may drive transitions in molecular state after repeated injury. Both single and repetitive TBIs have been associated with hyperphosphorylated tau and CTE-like phenotypes (65), and our shotgun proteomics revealed a marked shift in protein abundance patterns after three impacts (cumulative KE), consistent with a threshold-like transition in this model. The 24-hour inter-injury interval may further shape the response, given human second impact syndrome (66), and the possibility that incomplete recovery between impacts amplifies network-level dysregulation. Longitudinal profiling after five impacts revealed sex- and time-dependent remodeling of mitochondrial/metabolic pathways, oxidative stress responses, and calcium-dependent kinase activity, with the largest divergence occurring at discrete post-injury windows (notably 7 DPI for metabolism-enriched clusters) and persistent sex differences in calcium/kinase-associated proteins.

The enrichment of mitochondrial and redox-associated terms suggests that energy homeostasis is a major axis of post-rTBI remodeling. Proteomic modules enriched for oxidative phosphorylation and broader energy metabolism were most perturbed at 7 DPI, consistent with delayed metabolic compensation described in mammalian rTBI models (67). Oxidative stress pathways, including glutathione-related terms(68), were also enriched, consistent with established links between mitochondrial dysfunction, elevated ROS (69), and neurodegeneration (70). These observations highlight mitochondrial redox regulation as a mechanistic node that may contribute to sex-specific recovery dynamics (**Figure 9B**). In parallel, enrichment of calcium-dependent serine/threonine kinase activity (including CaMKII) and its sexually dimorphic trajectory is notable given known links between CaMKII signaling, apoptotic pathways, and motor impairment after TBI in rodents (71). The male-biased reduction in this module over time may reflect a sex-specific course of proteomic re-equilibration, consistent with prior reports of sexually dimorphic post-injury protein dynamics (72, 73).

To connect these pathway-level signals to candidate effectors, we examined representative proteins within modules associated with locomotion, redox regulation, and calcium signaling. Cluster 4, which included proteins annotated with locomotor behavior, contained cabeza (caz), Akap200, Lamin (Lam), and SERCA, suggesting a link between injury-associated proteomic remodeling and the locomotor phenotype. Notably, caz has been implicated in neurodegenerative disease, including frontotemporal dementia (74). In parallel, we observed reduced abundance of multiple glutathione S-transferase isoforms with increasing injury burden, consistent with diminished detoxification capacity and a shift toward oxidative stress vulnerability. Because glutathione-dependent redox regulation is central to ROS buffering (68), these patterns implicate glutathione pathway modulation as a plausible approach to neuroprotection; for example, glutathione reductase overexpression increases resistance to oxidative stress and extends longevity under hyperoxic conditions in flies (75).

Sex- and time-dependent differences were also prominent in Cluster 6, enriched for calcium-dependent serine/threonine kinase activity, including CaMKII. CaMKII is a key node linking calcium signaling to phosphorylation-dependent stress responses and apoptotic pathways, and it has been associated with motor impairment and apoptotic signaling in rodent TBI models, where CaMKII inhibition mitigates injury-associated deficits (71). The consistent decrease of this module in males across post-injury time points suggests a sex-specific trajectory of kinase-pathway remodeling that may contribute to divergent recovery dynamics.

Finally, manual inspection of 1 DPI protein sets highlighted early differential expression of locomotor- and mitochondria-associated proteins (e.g., caz, TTC19, and Plc21C) with sex-specific patterns (**Figure S6**). Plc21C (orthologous to human PLCB1) modulates calcium signaling and has been linked to neurodevelopmental and neurodegenerative phenotypes (76), while TTC19 is required for mitochondrial complex III function, and TTC19 disruption causes neurodegeneration in both humans and flies (77, 78). Together, these candidates provide a mechanistic bridge between the enriched GO categories (mitochondria/redox and calcium signaling) and the observed behavioral deficits following rTBI.

rTBI produced neurodegeneration at 14 DPI as evidenced by increased vacuolization and TUNEL-positive cells, supporting our model’s ability to capture structural sequelae beyond acute behavior. Because tau aggregation is pathognomonic of CTE (65), we also examined tau dependence. Although wild-type injured flies showed tau accumulation around vacuoles, preliminary data from a dTau knockout background indicate that vacuolization can still increase after rTBI in the absence of dTau, suggesting that tau accumulation is not strictly required for neurodegeneration in this setting. This aligns with prior evidence that endogenous dTau deletion does not necessarily impair baseline neuronal function (79) and is consistent with the possibility that tau accumulation may be downstream of injury-driven degeneration rather than its initiating cause. These findings motivate future work to test whether tau modulates specific molecular pathways (mitochondrial or immune responses) after rTBI, as suggested by transcriptional changes reported in dTau-deficient flies following TBI (80).

Behavioral recovery was not uniform across domains. While locomotor impairments largely resolved over time, both sexes exhibited persistent decision-making deficits in the VBFD assay through 21 DPI, indicating that cognitive-like outcomes may be more durable than gross locomotor readouts in this model. This dissociation is consistent with the likelihood that distinct neural substrates underlie locomotion versus decision-making in flies (central complex versus mushroom bodies) (81),(^82^). The presence of neurodegenerative changes at 14 DPI further supports the need to connect region-specific pathology to behavioral outcomes, including quantitative mapping of vacuolization to structures implicated in the VBFD phenotype. Although our model injures one fly at a time, limiting high-throughput injuries, it allows for rapid single-fly processing. It takes less than one minute to injure one fly, offering an innovative and highly reproducible approach to modeling rTBI in *Drosophila*. Behavioral assays capture specific dimensions of function and cannot fully recapitulate the spectrum of human symptoms. Future work should also refine injury timing (shorter/longer intervals), formally model change points in proteomic state across injury counts, and test candidate protective pathways suggested by the data (mitochondrial redox regulators such as SOD (83),(^84^)).

In summary, we present a reproducible *Drosophila* rTBI platform and demonstrate that repeated head injury produces long-lasting behavioral deficits, neurodegeneration, and dynamic proteomic remodeling. The sex-dependent timing of recovery, combined with mitochondrial/redox and calcium signaling signatures, provides a mechanistic framework for probing sex-specific vulnerability and resilience after repetitive head trauma.

## Materials and Methods

### RESOURCE AVAILABILITY

Lead contact

Further information and requests for resources should be directed to and will be fulfilled by the first author, Nicole Katchur.

Materials availability

This study did not generate new unique reagents.

Data and code availability

● All datasets generated during this study are available upon request.
● Custom R scripts used for locomotor behavior analysis are available upon request.
● Arduino scripts available on request.

### EXPERIMENTAL MODEL AND SUBJECT DETAILS

*Drosophila melanogaster* stocks and husbandry

Wild-type *Oregon-R* flies were obtained from J. Schottenfeld-Roames. *TauKO* flies (Bloomington stock #64782, w[*]; TI{w+mC]=TI}tau[KO], Donor: L. Partridge (Max Planck Institute)). Flies were maintained at 25°C on a 12 h light/dark cycle on standard cornmeal/agar media (Princeton University’s *Drosophila* Media Core). Newly eclosed flies were collected within a 12-hour window and separated by sex 48 h post-eclosion using brief CO₂ anesthesia. Experimental injuries were performed 3 days post-eclosion.

## METHOD DETAILS

### Assembly and calibration of the ballistic impactor

The ballistic impactor was adapted from Sun and Chen (2017) (27) with modifications (**Figure S2A**). A 1 mL tuberculin syringe barrel (without plunger) was cut at the 1 mL mark. A 200 μL pipette tip was modified by removing the aerosol barrier, which was repurposed as the impactor. A 1 mL syringe needle cap was inserted into the syringe barrel to stabilize the fly holder. The assembled apparatus was mounted vertically using a clamp and ring stand. CO₂ flow (1–10 psi) was controlled via a solenoid valve (LHDD0400100, The Lee Company) triggered by an Arduino UNO R3 microcontroller. Pressure calibration was performed using a digital manometer (DM8215, General). Kinetic energy (KE) was calculated from slow-motion video (iPhone 14 Pro, iOS 17.4) using KE = ½ × *m* × (*d/t*)², where *m* = mass of the impactor, *d* = distance traveled, and *t* = time to impact. Linear regression of KE vs. pressure showed R² = 0.933 (**Figure S2C**).

### Assembly of electronic control circuit

The solenoid valve was controlled by the Arduino UNO R3 interfaced with a solid-state relay (SSR) and powered by a 24 V DC supply (**Figure S2B**). Connections were made using 6” male-to-female jumper wires. The Arduino IDE (v2.3.2) controlled valve pulses (100 ms duration) and Processing 4 (v4.0.1) provided a graphical user interface (GUI).

### Fly brain injury protocols

**Manual Injury Device**: CO₂ flow was delivered using a FlyBuddy flow regulator at 3.0, 5.0, or 7.0 L/min.

**Microcontroller-Triggered Device**: Injuries were induced using CO₂ pulses at 1.0 ± 0.1, 5.0 ± 0.1, or 10.0 ± 0.1 psi. Flies received either one impact per day for five days (longitudinal protocol) or 1–5 impacts in one day (progressive protocol). Flies were loaded headfirst into modified pipette tips, ensuring proboscis protection. Post-injury, flies recovered in vials at 25°C and were transferred to fresh food every 2–3 days.

### Behavioral assays

#### Locomotor activity

Locomotion was quantified as in Iyengar et al. (2012) (42). Flies (n = 10 per group) were placed in 55 mm 1% agarose-lined Petri dishes to prevent flight and acclimated for 2 hours before recording in a behavior tent. Videos were captured using a GoPro Hero10 (2,048 px, 30 fps).

Videos were exported to an Apple MacBook Pro computer and converted from .mp4 files to .avi files using homebrew and ffmpeg packages in the Terminal. Video files were then opened and analyzed in the IowaFLITracker GUI using MATLAB. Distance and velocity files were exported as .matlab files and further analyzed by R Studio code (refer to Appendix A.3, A.4). In brief, the analysis using code in R removed video artifacts and trimmed the video to the first 18,000 frames (10 minutes). Average velocity was calculated by averaging the velocity of the fly when the fly was moving between 3 mm/s and 60 mm/s. The frames spent at rest were calculated by analyzing the total frames the fly’s speed was between 0 and 3 mm/s. To calculate the time at rest, the frames spent at rest were divided by 30 frames per second. Cumulative distance, average velocity, and time at rest for each fly was exported to .csv files and further analyzed for statistical significance in GraphPad Prism 10.

### Value-based feeding decision assay

Performed as in Yu et al. (2021)(52) with minor modifications. Briefly, flies were food-deprived for 8 hours and placed in Petri dishes containing droplets of 150 μM sucrose (nutritious) and 150 μM arabinose (non-nutritious), dyed with 0.01% erioglaucine or 0.1% Food Red No. 106. Choice was scored under a stereomicroscope after 2 hours.

### Histology and imaging

Flies subjected to repeated head trauma were collected 1 day, 14 days or 28 days following the last injury. Flies were embedded in paraffin and brain vacuoles were assessed using the protocol outlined by Sheng et al. 2023 (87). In brief, flies were rapidly decapitated under mild CO_2_ and then fixed in Bouin’s fixative (Sigma-Aldrich) for five days inside of Eppendorf microtubes. Samples were transferred to lens paper inside of Tissue Path Cassettes (FisherScientific, Hampton, NH). Cassettes were transferred into leaching buffer (50 mM Tris, pH 8, 150 mM NaCl) for 24 hours at room temperature, and the solution was changed every 8-12 hours until no longer yellow. Cassettes were then transferred to 70% EtOH for dehydration and processed by the HistoCore Tissue Processor (Leica, Deer Park, IL). Heads were blocked in paraffin and sectioned into 8µm thick ribbons. Ribbons were deparaffinized by washes in Histoclear I and II (National Diagnostics, Beachwood, OH), and stained using a standard H&E staining protocol, TUNEL HRP-DAB kit (ab206386, abcam, Cambridge, UK) according to the manufacturer’s instructions, or with a primary antibody. Sections were mounted with Permount (FisherScientific, Hampton, NH) to assess brain vacuolization. Sections were imaged on a Nikon A1 confocal microscope under 20× objective, 1× magnification with fixed exposure settings. Vacuolization was assessed using FIJI (ImageJ 1.0) by thresholding images (to make binary) and using the built-in function “Analyze Particles” to measure vacuoles. To minimize the inclusion of tissue artifacts from the processing procedure, vacuoles with circularity of greater than 70% and an area greater than 30µm^2^ were measured. Section matching of serial sections among the groups was performed, and the total vacuolization area for each brain was calculated by summing the vacuolization areas for each section. The total area of each section was also determined, and the percent vacuolization was calculated by the total area of vacuolization across the three sections over the total tissue area of the three sections per brain.

To assess dTau accumulation, slides containing rTBI dTauKO brain sections or Oregon-R wild-type rTBI brain sections were blocked in 5% milk and 10% NGS in TBS-T for 1 hr. The slides were then incubated in rabbit anti-dTau custom antibody (ABClonal) in a 1:2000 dilution for 24 hrs. After primary incubation, slides were incubated with a secondary antibody mouse anti-rabbit HRP (1:5000, Jackson ImmunoResearch) for 2 hrs. Slides were developed using the HIGHDEF blue IHC chromogen kit (Enzo Life Sciences) for 4.5 min. Slides air-dried, were coverslipped, and imaged as outlined above.

### Proteomic Analysis

#### Sample Preparation

Following rTBI, 50 flies per condition were placed on dry ice, and heads were dissected using Dumont Biology tweezers (Ted Pella). Heads were transferred to 1.5 mL tubes with 5% SDS and 50 mM TEAB (pH 7.1), homogenized using a motorized pestle, and centrifuged at 10,000 rpm for 1 min. Supernatants were transferred to fresh tubes. Protein concentrations were determined using the Pierce BCA assay (#23225, Thermo Scientific), and readings were taken on a NanoDropOne. HALT Protease/Phosphatase Inhibitor (#78440, ThermoFisher) was added to 1X final concentration.

### Mass Spectrometry

Samples were processed with S-TRAP micro spin columns (ProtiFi) per the manufacturer’s instructions. Tryptic peptides were dried in a SpeedVac and resuspended in 6 μL of 0.1% formic acid (pH 3). 2 μL (∼360 ng) was injected per run into an Easy-nLC 1200 system. Peptides were loaded onto a 45 cm × 100 μm ID nano-column packed with 1.9 μm C18-AQ resin (Dr. Maisch) at 50°C using a 2-hour gradient at 360 nL/min. MS1 spectra were acquired on an Orbitrap Fusion Lumos in positive mode (120,000 resolution, 375–1500 m/z, AGC target 4e5, 54 ms injection time). MS2 spectra were collected in the ion trap with 35% HCD energy. Dynamic exclusion was set to 60 s with a 3 s maximum cycle time. Peptides were isolated using a 1.6 m/z quadrupole window. FAIMS was used with a DV of 5000V and CVs of −50, −63, and −81V.

### Protein Identification

Raw files were analyzed using Sequest HT in Proteome Discoverer 2.5 with the *Drosophila melanogaster* UniProt database (UP000000803). Search parameters: 10 ppm MS1 tolerance, 0.6 Da MS2 tolerance, up to 2 missed cleavages. Fixed modification: carbamidomethylation (C).

Variable modifications: oxidation (M), phosphorylation (S/T/Y), N-terminal acetylation. Peptide/protein validation was performed in Scaffold v5.3.1 using the Local FDR algorithm (>95% peptide, >99.9% protein probability, ≥2 unique peptides). Protein probabilities were assigned using Protein Prophet. Total ion current was normalized, and values were exported for analysis.

### Manual clustering of the proteomic data

Normalized, log₂-transformed fold changes (TBI vs. sex-matched control) were used to assign proteins into nine expression cohorts. A threshold of ±2 log₂ fold-change (4-fold change) defined differential abundance. Cohorts reflected sex-specific expression patterns: increased, decreased, or unchanged in males vs. females, resolving sexually dimorphic responses that may be masked in bulk analysis.

## QUANTIFICATION AND STATISTICAL ANALYSIS

All statistical tests were two-tailed unless otherwise indicated, with a significance threshold of *p* < 0.05. Data are presented as mean ± SD unless otherwise noted in the figure legends (mean ± SEM for select histology readouts). Sample sizes (*n*) and the specific statistical test applied for each experiment are reported in the corresponding figure legends.

### Apparatus Calibration

For impactor calibration, the relationship between pressure setting and derived kinetic energy was assessed by linear regression, with goodness-of-fit reported as R²

### Survival Analysis

Survival was analyzed using Kaplan–Meier survival curves with group comparisons performed using log-rank tests (Mantel–Cox). Dose/intensity-response effects across injury conditions were assessed using a log-rank test for trend, and multiplicity was controlled using Holm–Šídák multiple-comparisons correction where applicable. When hazard ratios (HR) are reported, they reflect relative mortality risk estimated from proportional hazards modeling.

### Locomotion Assay

For acute post-injury locomotor outcomes (e.g., cumulative distance at 1 day post-injury), injured and control groups were compared within sex using a two-tailed Mann–Whitney U test, and a two-way ANOVA was used to evaluate sex-by-injury interaction effects where indicated. For injury-threshold experiments (1–5 injuries) and longitudinal time-course analyses (1, 7, 14, and 21 days post-injury), outcomes were analyzed using two-way ANOVA with Tukey’s multiple-comparisons test (performed as specified per figure/sex), with multiplicity-adjusted post hoc comparisons used to localize significant effects. Rest time was analyzed as a continuous outcome derived from the locomotor tracking pipeline (rest defined as movement below 3 mm/s). To test whether injury effects on locomotion differed by sex over time, a factorial model including Sex, Injury, and Time was evaluated, including the Sex × Injury × Time interaction; multiplicity-adjusted contrasts were used for planned post hoc localization of interaction effects. For the prespecified sex comparison at 14 days post-injury after normalization to sex-matched controls, a Welch’s two-sample *t* test was used with Holm–Bonferroni adjustment; effect size was quantified using Hedges’ *g*. The analysis was performed using code in MATLAB2025b.

### Value-based Feeding

Feeding preference outcomes were quantified as the percentage of flies choosing sucrose out of the total tested, including non-choosers. Preference outcomes across time and injury condition were analyzed separately by sex using two-way ANOVA with Tukey’s multiple-comparisons test; replicate structure and sample sizes are provided in the figure legends. Related secondary outcomes (arabinose choice and “no preference”) were analyzed with the same two-way ANOVA/Tukey framework.

### Histology

For brain vacuolization and related histologic quantifications, injured and control groups were compared using unpaired two-tailed *t* tests, with the biological replicate defined at the level specified in the figure legends (e.g., sections nested within brains, summarized per brain as reported). For dTauKO analyses, group differences in vacuolization were likewise assessed using unpaired two-tailed *t* tests.

### Proteomic Quantification

Proteomic heatmaps display relative abundance changes (fold-changes or injured-to-control ratios as indicated), with clustering performed using k-means (k = 6). For longitudinal analyses and supplemental manual clustering, protein abundance ratios (injured vs age-matched controls) were log₂-transformed prior to clustering/visualization. Gene ontology enrichment analyses were conducted using ShinyGO (v0.80), with fold enrichment and false discovery rate (FDR) values reported.

## POWER ANALYSIS

Sample sizes were calculated to achieve 80% power with α = 0.01, based on prior studies (27).

## Acknowledgments

We thank G. Broussard for circuit design advice and J. Schottenfeld-Roames for providing *Oregon-R* flies. We thank William dePaola for his contribution to the locomotion assay and his engagement with the research process.

## Author Contributions

N.J.K. designed research; performed research; analyzed data; and wrote the paper. J.Y. performed research; contributed new reagents/analytic tools; analyzed data; and contributed to writing and revising the paper. L.S. contributed to research design and planning. H.S. performed research and contributed to research design and planning. D.A.N. designed research; supervised and mentored the project; and contributed to writing and revising the paper.

