## Supplemental Text and Figures for "Modeling Sex Differences and Neurodegeneration in Repetitive Traumatic Brain Injury Using *Drosophila*"

**Supporting Information Text**

Despite the heterogeneity of TBI in humans, specific areas of the brain are more vulnerable to mechanical stress. Deeper cortical structures, such as the hippocampus, are most susceptible to rotational head injury (85, 86). To determine which areas of the fly brain are most vulnerable to rTBI, we categorized the *Drosophila* brain anatomy into three major regions, the neuropil, the lobula, and the medulla of the optic lobe (**Figure S5A**). These regions were chosen because they could be reliably distinguished in each of the brain sections. Within controls, 58% of the observed vacuoles were present in the neuropil region, while 25% were present in the medulla, and 17% were present in the lobula (**Figure S5B**). In comparison, 36% of vacuoles were observed in the neuropil region, while 32% were located in the lobula and 28% in the medulla (**Figure S5B**). This shift in vacuole distribution suggests that the lobula region may be particularly susceptible to the impact of rTBIs, though additional studies will be needed to reach significance.

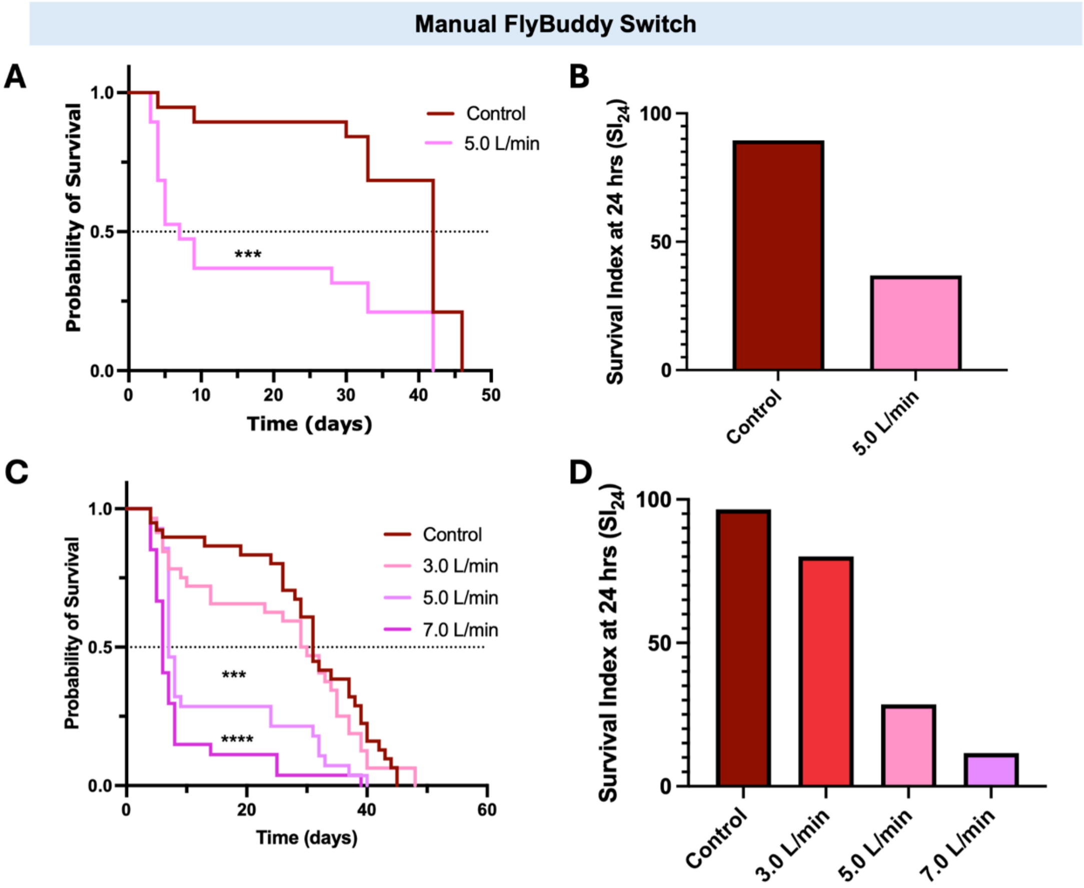


**Figure S1. Survival effects of repetitive impacts using the manual-switch FlyBuddy apparatus.** (A, C) Wild-type (*Oregon-R*) males showed significant decreases in survival probability after five repetitive traumatic brain injuries (rTBIs) at $CO_2$ flow rates of 5.0 L/min and 7.0 L/min, respectively. (B, D) The 24-hour survival index, defined as the percentage of flies alive 24 hours post-injury, decreased with increasing $CO_2$ flow rate. Injuries were delivered using the manual-switch FlyBuddy $CO_2$ flow regulator. Data are presented as mean ± SD. ***$p < 0.001$, ****$p < 0.0001$; Log-rank (Mantel-Cox) test and Log-rank trend test with Holm-Šídák's multiple comparisons correction. $n$ = 19 flies per group (A), $n$ = 26–30 flies per group (C).

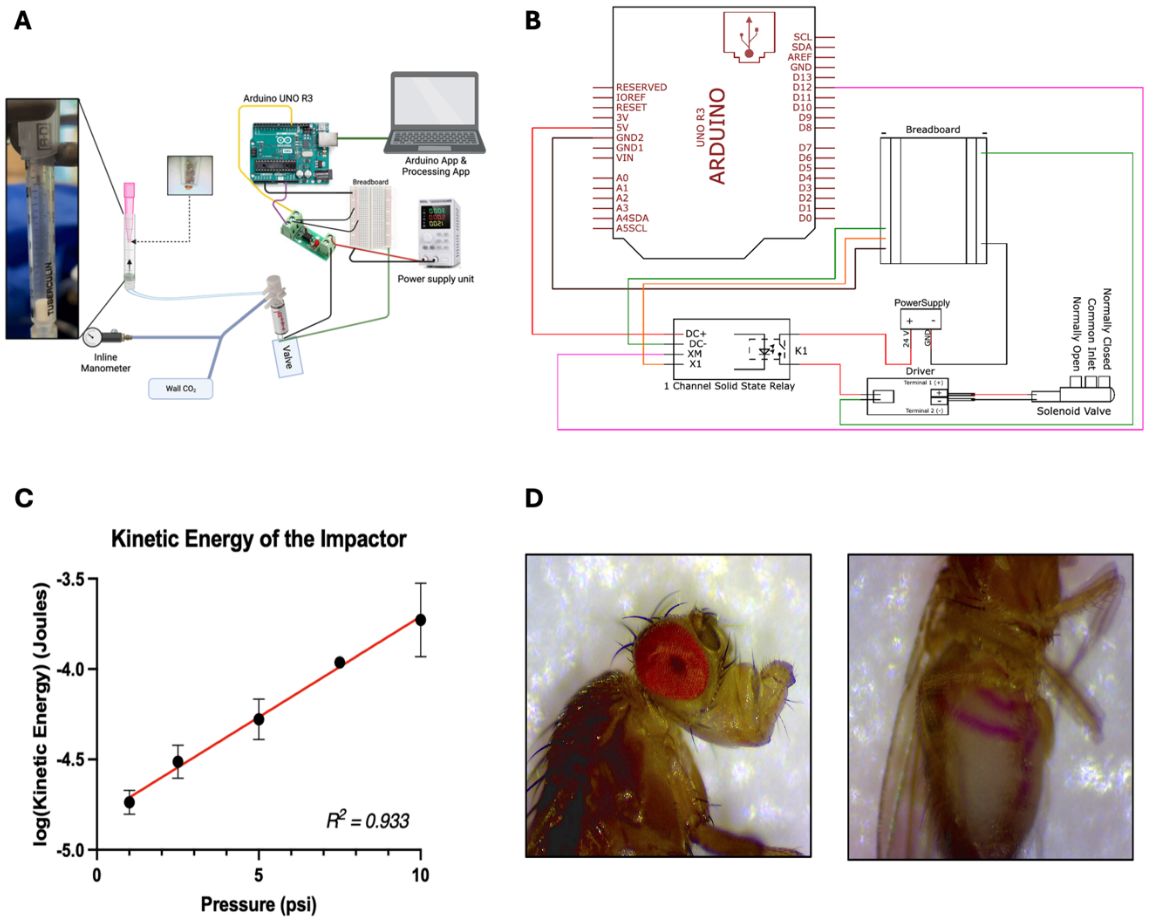


**Figure S2. Design and calibration of an improved apparatus for delivering rTBI in *Drosophila*.** (A) Schematic of the device showing a fly positioned in a 200 µL pipette tip with its proboscis tucked to prevent injury. A $CO_2$ pulse (1.0 ± 0.1 psi) propels a pipette plug impactor through a syringe barrel to strike the fly's head. Flow is regulated by a solenoid valve triggered by an Arduino Uno R3 microcontroller (100 ms pulse). Calibration was performed across 1–10 psi. Image created with BioRender.com. (B) Circuit diagram showing Arduino and solid-state relay (SSR) connections. Diagram created using EasyEDA Std (Guangdong, China). (C) Kinetic energy (KE) of the impactor was calculated during calibration using $KE = 0.5 \times m \times (d/t)^2$, where $m$ is mass, $d$ is distance traveled, and $t$ is time. Data are mean ± SD; linear regression goodness of fit: $R^2 = 0.933$. $n$ = 3 trials per pressure setting. (D) Representative images showing (left) no visible external head damage following rTBI and (right) intact intestinal barrier assessed with sucrose and red dye after five impacts. Image created with BioRender.com.

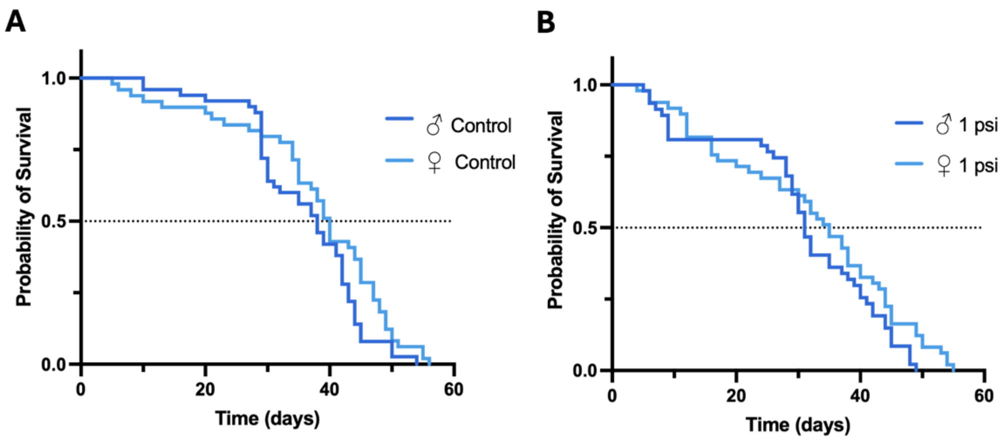


**Figure S3. Survival effects of repetitive impacts in males and females at 1 psi.** (A, B) No significant differences in survival probability were observed between control and injured groups in wild-type (*Oregon-R*) males and females subjected to five impacts at 1 psi. Data are shown as mean ± SD. Log-rank test. *n* = 46–50 flies per group.

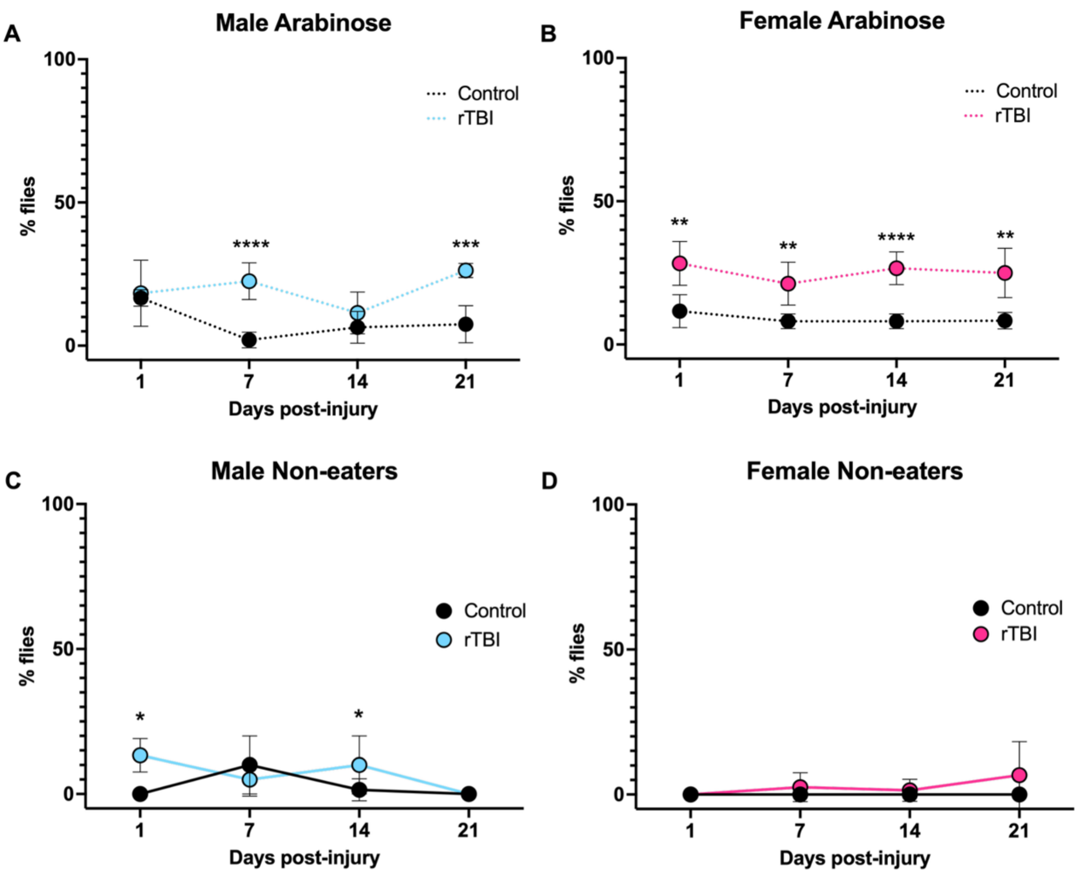


**Figure S4. Feeding decisions of flies that did not choose sucrose following rTBI.** (A, B) Percentage of flies choosing arabinose over time in males and females, respectively. (C, D) Percentage of flies showing no preference for sucrose or arabinose over time in males and females, respectively. Data are presented as mean ± SD. *$p < 0.05$, **$p < 0.01$, ***$p < 0.001$, ****$p < 0.0001$; two-way ANOVA with Tukey's multiple comparisons test. $n = 10$ flies per group; 3–7 groups per time point.

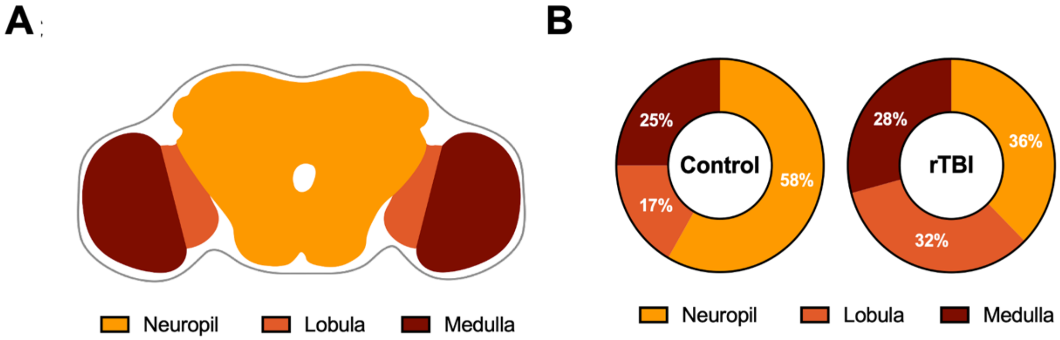


**Figure S5. Regional distribution of brain vacuoles following rTBI.** (A) Diagram showing the three brain regions analyzed for vacuole localization. (B) Percentage of total vacuoles detected in each brain region for control and rTBI flies. *n* = 12–25 vacuoles per group.

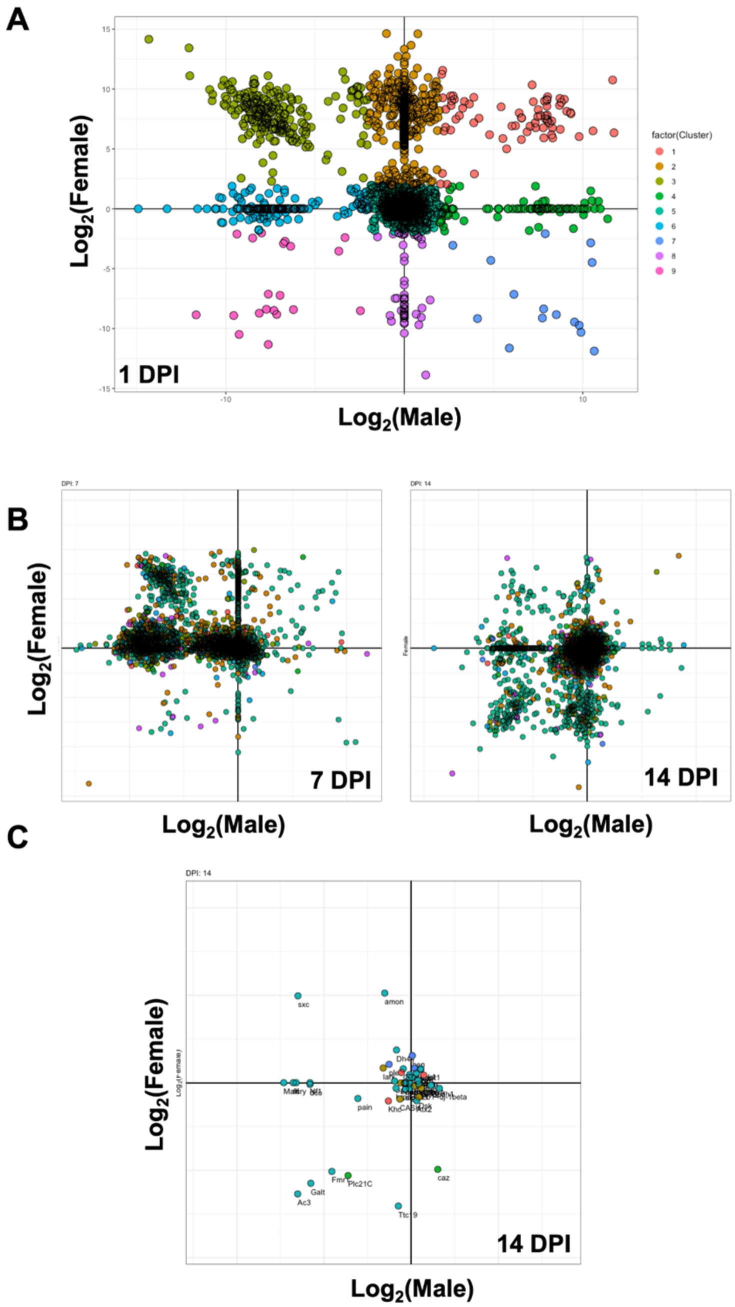


**Figure S6. Manual clustering of longitudinal proteomic changes post-rTBI.** (A) Manual clustering of protein abundance in males and females identified nine distinct clusters. Protein abundance ratios (injured vs. age-matched controls) were $\log_2$-transformed. (B) Representation of the same clusters at 7 and 14 days post-injury. (C) Enriched gene ontology (GO) terms for locomotor behavior at 14 days post-injury, with colors corresponding to cluster assignments from one day post-injury.

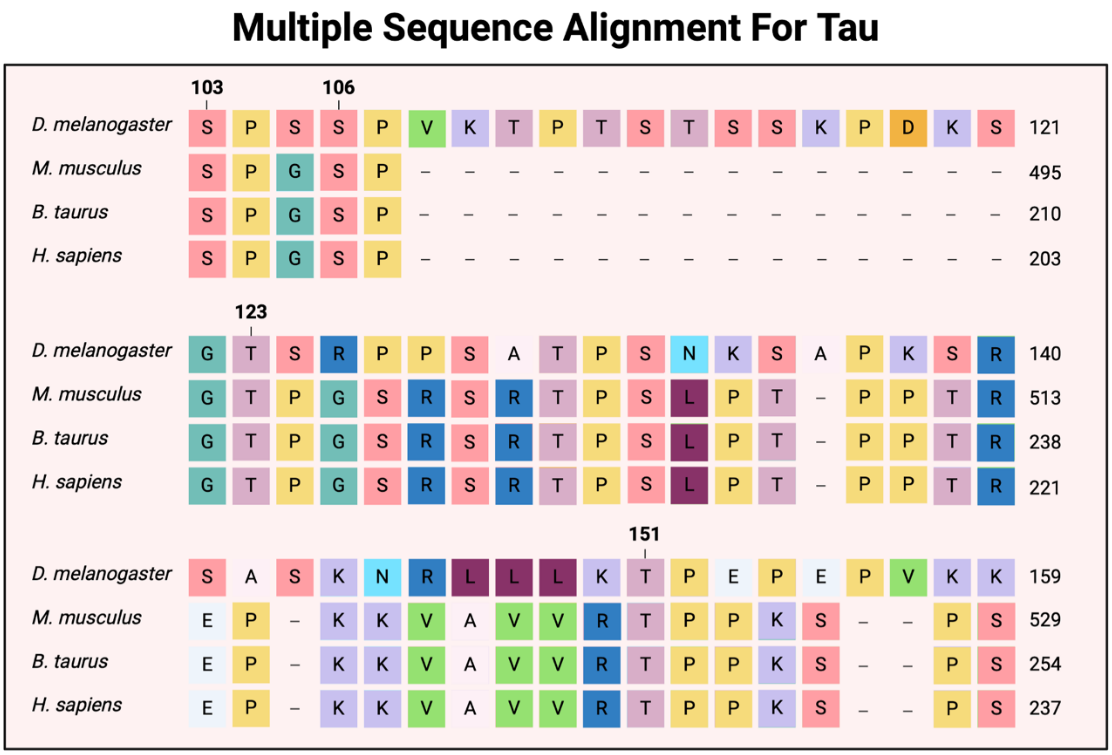


**Figure S7. Sequence alignment of tau proteins across species.** Alignment of tau protein sequences from *Mus musculus* (mouse), *Bos taurus*(cow), *Homo sapiens* 4-repeat tau isoform (human), and *Drosophila melanogaster* (fruit fly). Conserved amino acids implicated in chronic traumatic encephalopathy pathology are bolded in the *Drosophila* sequence. Alignment performed using UniProt (The UniProt Consortium, 2023. Nucleic Acids Res. 51:D523–D531). Image created with BioRender.com.
